# Phosphorylated α-Synuclein shapes changes in brain structure, task-related activity and functional connectivity in M83 mice

**DOI:** 10.64898/2026.09.16.752140

**Authors:** Amr Eed, Vladislav Novikov, Shahrzad Bahrampour, Kelly Summers, Stephanie Tullo, M. Mallar Chakravarty, Wen Luo, Irina Shlaifer, Thomas M. Durcan, Vania F. Prado, Ali R. Khan, Marco A.M. Prado, Ravi S. Menon

## Abstract

Phosphorylated α-synuclein at S129 (pS129) is the predominant form of α-synuclein in the aggregates known as Lewy bodies (LB) and Lewy neurites (LN), the pathological hallmarks of Parkinson’s disease (PD) and dementia with Lewy bodies (DLB), collectively known as synucleinopathies. The toxic nature of these inclusions remains contentious. To improve our understanding of these aggregates and their role in the structural and functional changes observed in synucleinopathies, we combined non-invasive structural and functional MRI in awake head-fixed M83 mice with whole-brain light-sheet fluorescence microscopy (LSFM) in the same animals following a unilateral injection of either human preformed fibrils of α-synuclein (PFF) or PBS into the dorsal striatum. We developed a multimodal framework integrating the non-invasive, mesoscopic nature of MRI with the cellular-level details of LSFM to further our understanding of the underlying pathology of synucleinopathies. Using this framework, we show that four months after injection, animals injected with PFFs showed ubiquitous spread of pS129 pathology throughout the whole brain and spinal cord consistent with anatomical connectivity from the seeding region, with some structures such as the midbrain more susceptible to pS129 accumulation. Compared to controls, PFF-injected animals showed widespread atrophy as well as a decrease in functional connectivity, while evoked activity in the somatosensory/somatomotor areas in response to spontaneous paw movements showed significant hypoactivation. These cellular, structural and functional changes were significantly associated with the proportion of pS129 accumulation determined in the same animals. Our results demonstrate the deleterious effects of pS129 accumulation and argue in favor of a toxic role played by these aggregates.

**One Sentence Summary:** Awake-MRI and whole-brain light-sheet microscopy reveal pS129 involvement in brain structure, task-related activity, and functional connectivity.

## INTRODUCTION

Aggregates of phosphorylated α-synuclein at S129 (pS129) are the predominant component of inclusions known as Lewy bodies (LBs) and Lewy neurites (LNs), the defining hallmarks of synucleinopathies such as Parkinson’s disease (PD) and dementia with Lewy bodies (DLB) (*1–3*). The formation of these inclusions is closely correlated with impairments in synaptic function that culminate in neuronal cell death in affected brain regions (*4–6*). In the absence of specific antibodies for aggregated α-synuclein, pS129 has become the de facto marker of pathological inclusions in synucleinopathies (*7*). Under normal physiological conditions, less than 4% of α-synuclein exists as pS129; this increases dramatically to over 90% within LB and LN deposits (*2, 3*).

However, whether pS129 enhances the toxicity of α-synuclein or plays a protective role remains an active area of debate, with studies lending support to both perspectives depending on context. Studies conducted in rodent models have shown that, while the phosphomimetic S129D α-synuclein is more prone to aggregation, the resulting aggregates are not inherently toxic and may confer a neuroprotective effect on dopaminergic neurons (*8*). Similarly, Ghanem et al. reported that S129 phosphorylation is a late event relative to aggregation and that it decreases the aggregation propensity and cytotoxicity of α-synuclein (*9*). Others have attributed the pS129 toxicity to the kinase mediating the phosphorylation, suggesting that it is the kinase rather than the phosphorylation itself that governs the toxicity or the neuroprotective effect of pS129 *(reviewed in 10)*.

Conversely, a study in Drosophila models of PD showed that S129 phosphorylation enhances α-synuclein neurotoxicity and leads to dopaminergic neuron loss, while blocking phosphorylation at S129 supressed dopaminergic neuronal loss while leading to an increase in inclusion formation (*11*). Furthermore, a recent study showed that excess pS129 has a dramatic effect on synaptic vesicle trafficking, leading to synaptic vesicle declustering and depletion (*5*).

Given its non-invasive nature, especially valuable for longitudinal studies, the use of structural and functional magnetic resonance imaging (MRI/fMRI) has gained considerable traction in both the diagnosis and the characterization of PD (*12–15*). Various studies have shown abnormal functional connectivity (FC) in PD and related disorders. Striatal networks appear to be the most affected, with abnormal connectivity with both the cortical circuits (*16*) and the brainstem (*17*). On the structural side, studies have shown volumetric differences in structures including the olfactory bulb and the striatum, as well as the substantia nigra, albeit with varying results in the latter *(see 15 for a review)*. Additionally, surface-based analyses have revealed cortical thinning (*18, 19*). MRI has also been employed to characterize mouse models of synucleinopathy, though such work remains limited (*20–24*). While functional MRI studies are sparse, structural MRI studies have demonstrated atrophy in various regions especially the striatum, thalamus, and brainstem akin to patterns observed in human studies (*20, 23*).

Although MRI can capture functional and structural changes at mesoscopic and macroscopic scales *in vivo*, the low-resolution nature of the images limits the ability to resolve fine-grained anatomical detail. Histological techniques offer superior spatial resolution but are inherently limited by their two-dimensional nature, which constrains the ability to capture the spatial complexity of biological tissues. Advances in volumetric microscopy, particularly light-sheet fluorescence microscopy (LSFM) and tissue-clearing techniques, have begun to bridge that gap, enabling whole-brain imaging at unprecedented detail while preserving the three-dimensional context of the sample (*25, 26*). Ultimately, combining such high-resolution three-dimensional imaging with the longitudinal power of MRI could produce important insights into neurodegenerative processes.

## RESULTS

We combined awake structural and functional MRI with whole-brain LSFM imaging to map the propagation of pS129 pathology in the mouse brain following unilateral α-synuclein PFF injection and to investigate how accumulation of pS129 alters brain structure and function. We also developed analysis pipelines that process raw LSFM datasets and produce quantifiable metrics in standard space, enabling correlations with outputs from the different MRI modalities. An overview of the experimental design and pipelines used is provided in Figure 1.

**Fig. 1.**
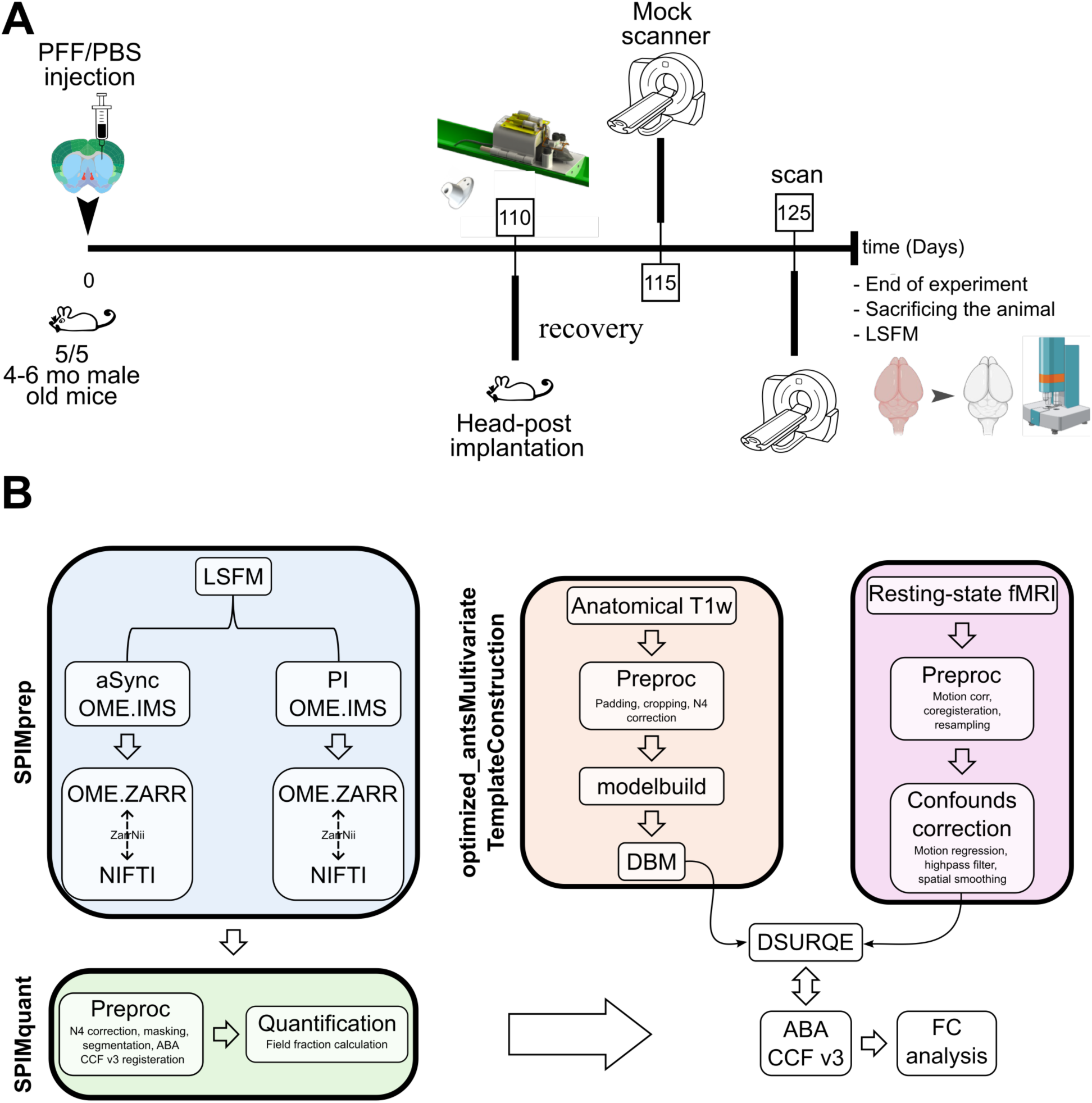
Overview of the experimental design and analysis pipelines. **A)** Experimental timeline showing either PFF or PBS injections in the dorsal striatum, headpost implantation, imaging, and LSFM. **B)** Analysis pipelines for processing of LSFM, anatomical, and functional data and aligning them to the same anatomical space to enable joint inferences. **Abbreviations:** ABA-CCFv3, Allen Mouse Brain Common Coordinate Framework v3; aSync, α-synuclein; DBM, deformation-based morphometry; FC, functional connectivity; LSFM, light-sheet florescence microscopy; mo, month; PBS, phosphate-buffered saline; PFF, preformed fibrils; PI, propidium iodide

### PS129 pathology propagation across the brain and the spinal cord

Whole-brain light sheet imaging revealed widespread pS129 pathology throughout the brain following inoculation of the α-synuclein PFF into the right dorsal striatum (Fig 2, Suppl. Fig 6, Suppl. video 1) compared to control animals injected with PBS. Field fraction, defined as proportion of pS129 positive voxels within each brain parcel, was used to quantify regional pathological burden. A low signal was observed in control animals injected with PBS across multiple structures, likely reflecting edge effects, and was substantially and consistently surpassed by animals injected with PFF in all affected structures.

**Fig. 2.**
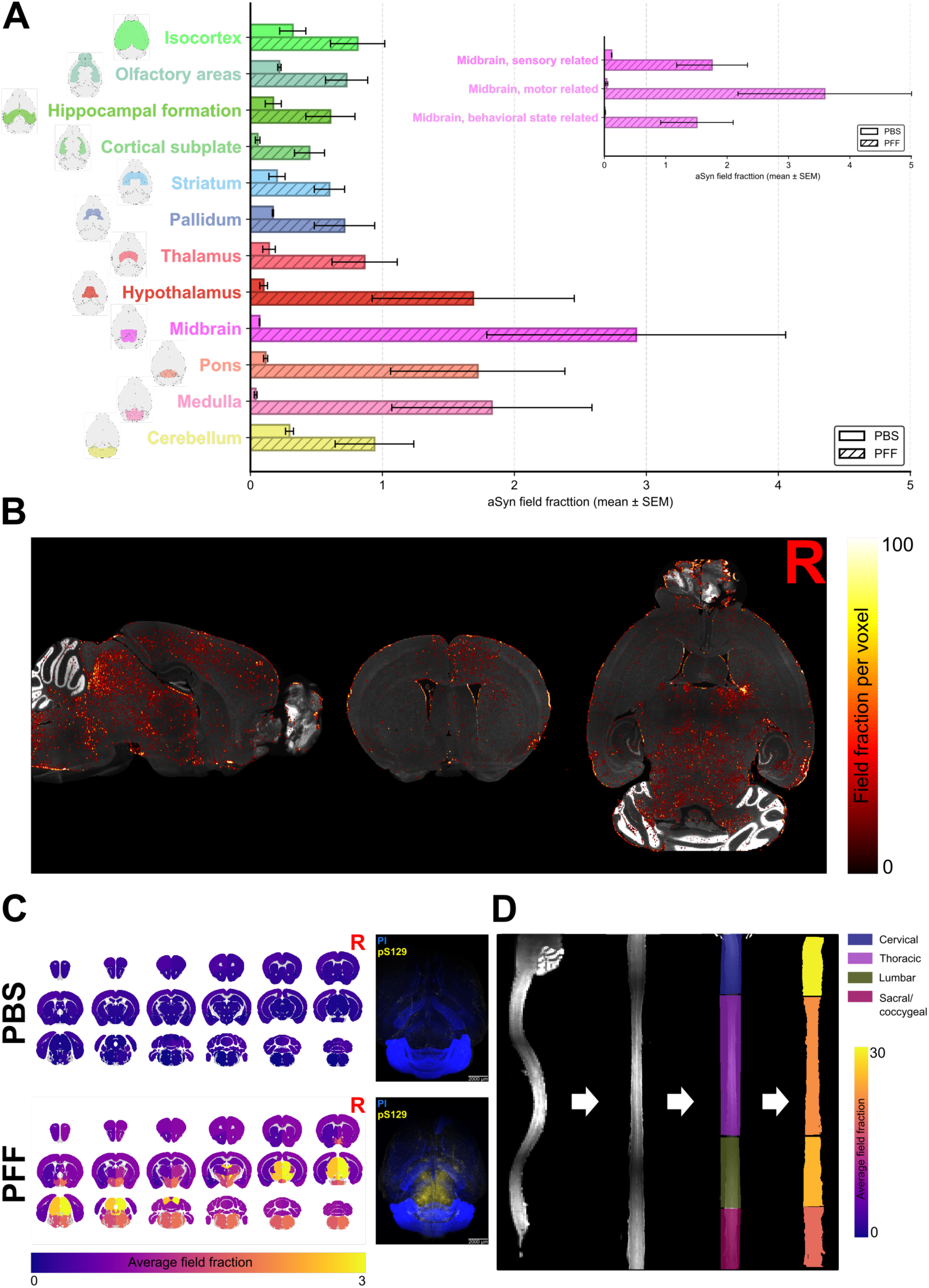
PS129 distribution across the brain and the spinal cord. **A)** Quantification of pS129 across major brain structures alongside its regional distribution within the midbrain substructures (bars, mean ± SEM). **B)** Representative pS129 field-fraction map from a PFF-injected animal overlaid on a PI background. **C)** Left, heat map of mean pS129 field fraction per ABA CCFv3 parcellation across all subjects; right, pS129 immunostaining from a representative subject. Scale bar: 2000 µm **D)** Spinal cord straightening and heat map of pS129 average field fraction from all subjects across the segments of the spinal cord. **Abbreviations:** ABA-CCFv3, Allen Mouse Brain Common Coordinate Framework v3; aSync, α-synuclein; LSFM, light-sheet fluorescence microscopy; PBS, phosphate-buffered saline; PFF, preformed fibrils; PI, propidium iodide; pS129, phosphorylated-s129-α-synuclein; R, right hemisphere; SEM, standard error of the mean.

Field fraction comparisons between ipsi- and contra-lateral hemispheres failed to reveal any differences that survived multiple comparisons correction (Suppl. Fig. 3); therefore, the data from both hemispheres were pooled for subsequent analysis.

Although the right dorsal striatum was the pathology epicenter, the pS129 pathology spread to structures throughout the brain. At the coarse parcellation level (Suppl. Fig. 1), the midbrain showed the highest pS129 burden across the brain, followed by hindbrain structures (the pons and medulla) and interbrain structures (particularly the hypothalamus). The cerebellum also exhibited substantial pathology, as did the olfactory areas, pallidum, and striatum. Notably, cortical and limbic structures including the isocortex and hippocampal formation showed considerable pathology as well.

Fine-grained parcellation revealed a more nuanced pattern of pathology (Suppl. Fig. 6). Within the midbrain, the motor-related areas, encompassing the substantia nigra and the ventral tegmental area, appeared to be the subdivision with the most pathology across in the entire brain, harboring the highest pS129 burden of any region. Across the midbrain and hindbrain regions, motor-related subdivisions consistently showed higher pathology burden than their sensory-related and behavioral state-related counterparts (Fig. 2A). The lateral zone of the hypothalamus exhibited the highest burden among hypothalamic regions.

The pathology appears to spread along the rostrocaudal axis, with caudal spread being more pronounced (Fig. 2A&C). Spinal cord samples showed a rostrocaudal gradient as well, with the cervical and thoracic regions exhibiting more pathology than the lumbar and sacral/coccygeal regions (Fig. 2D, Suppl. video2). High inter-animal variability was observed in both parcellations, particularly in the most affected areas, as reflected by the relatively large error bars. The variability is likely stemming from the heterogeneous nature of the misfolding and spread of pS129, as all the experimental variables (injection site, injected dose, and PFF batch used) were consistently maintained among experiments.

### PFF-injection induces bimodal structural changes

Unilateral PFF injection in the dorsal striatum induces pervasive neuroanatomical changes spanning cortical, subcortical, and cerebellar structures (Fig. 3). Voxelwise comparison of the relative Jacobian determinant maps from the DBM analysis between both groups yielded multiple significant changes.

**Fig 3.**
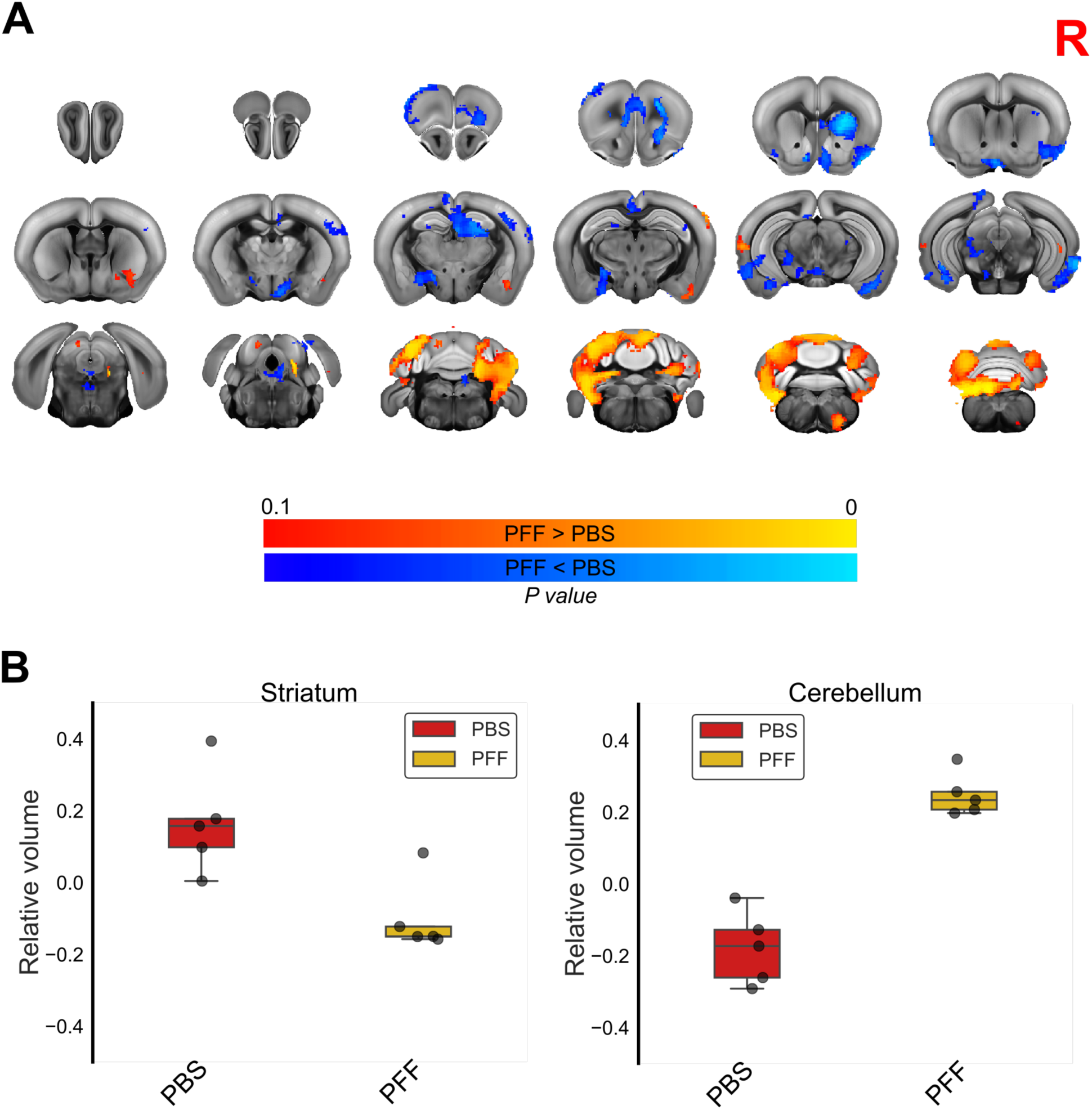
PFF injection induces structural changes. **A)** Voxelwise statistically significant differences between PFF and PBS-injected animals, where blue–light blue indicates local shrinkage and red-yellow indicates expansion in PFF-injected animals relative to controls. The statistical map is overlaid on the ABA-CCFv3 anatomical template. **B)** Log-Jacobian values extracted from peak voxels within the striatum (left) and cerebellum (right). **Abbreviations:** ABA-CCFv3, Allen Mouse Brain Common Coordinate Framework v3; DBM, deformation-based morphometry; PBS, phosphate-buffered saline; PFF, preformed fibril.

Similar to findings reported by Tullo et al. (*20, 24*), these changes were bidirectional, with different regions showing shrinkage or expansion across the brain. At the injection site in the right dorsal striatum, PFF-injected animals showed shrinkage relative to PBS-injected animals. The nucleus accumbens of the ventral striatum was similarly affected. This shrinkage extended to key isocortical regions such as the orbital, the somatosensory, the somatomotor area, and prelimbic areas showed changes, albeit mostly limited to the ipsilateral regions (Fig. 3). Expansion as well, though less extensive than contraction, was evident in the isocortex, most notably in posterior regions including the auditory and the temporal association areas. Bilateral contraction in the PFF-injected animals was observed in the olfactory regions such as the piriform area.

Ipsilateral volume reduction was also observed in Ammon’s horn regions of the hippocampus, additionally the dentate gyrus showed bilateral shrinkage. Multiple nuclei in the pallidum and the hypothalamus showed contraction on both sides of the brain. Interestingly, the amygdala showed volume reduction in both hemispheres.

Within the midbrain, the changes in the motor-related areas were more pronounced, the substantia nigra along with the ventral tegmental area contracted on the contralateral side; a similar trend was observed ipsilaterally but did not survive statistical thresholding. Within the hindbrain, the motor-related areas of the pons showed similar effect as well. The cerebellum showed a distinct pattern of expansion in both the cerebellar cortex and the cerebellar nuclei.

### Synucleinopathy disrupts functional organization

To assess whether the PFF injection might alter functional network organization within the brain, we first investigated the seed-based FC between the injection site in the right dorsal striatum and the rest of the brain. The AMBCA (*27*) was used to delineate the seed region overlapping with the injection location (Fig. 4A; see Methods). The PBS-injected animals showed strong bilateral FC between the seed and widespread cortical regions, including the anterior cingulate, prelimbic, somatomotor, and somatosensory areas, as well as the main olfactory bulb (Fig. 4B). The PFF-injected animals, by contrast, showed weaker bilateral connectivity. Subcortically, the seed area in the PBS-injected animals had stronger correlation with the contralateral striatum and the hippocampus compared with PFF-seeded group (Suppl. Fig. 7).

**Fig. 4.**
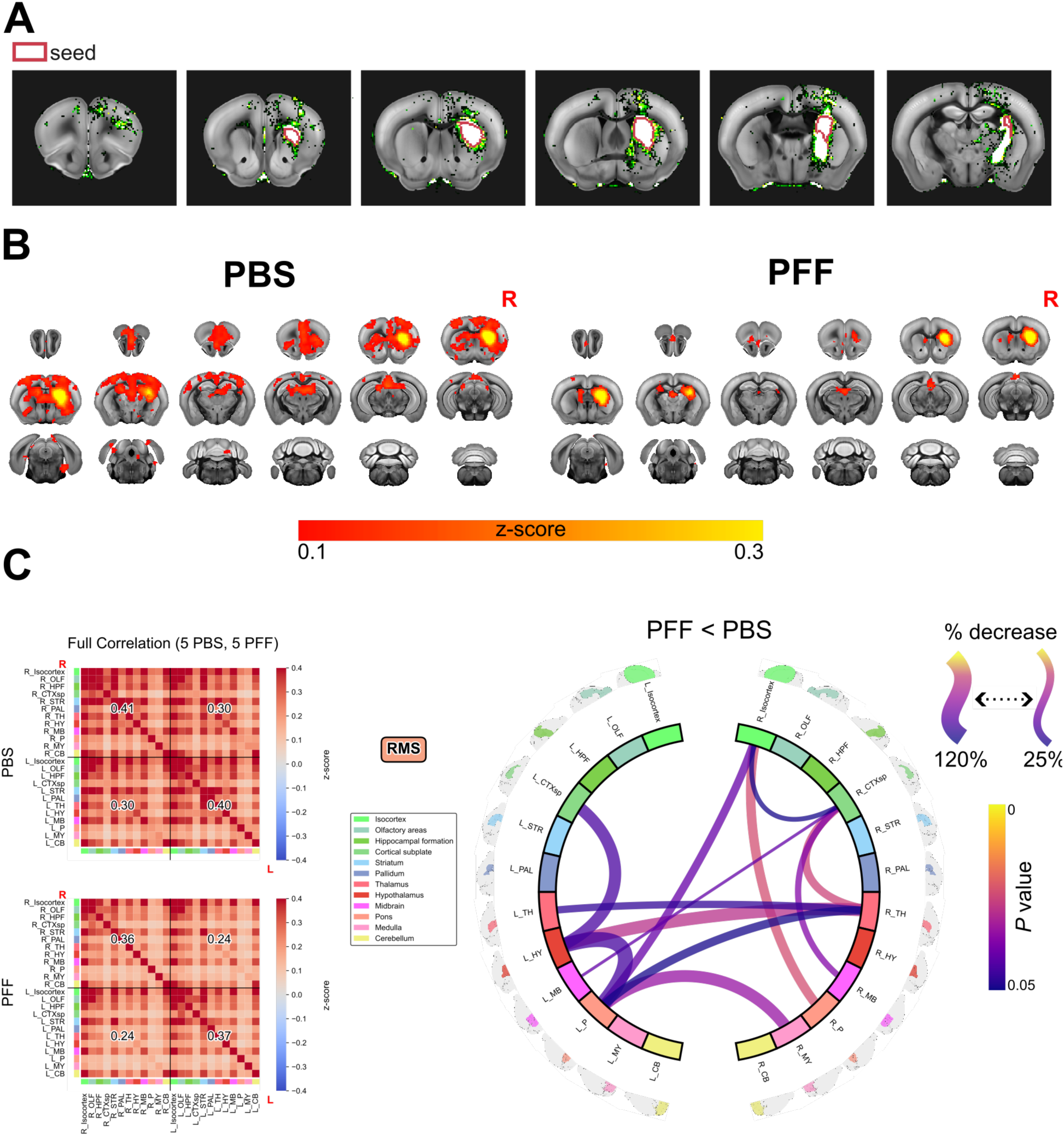
PFF injection disrupts functional organization. **A)** Maximal projection density from injection site at the dorsal striatum, showing the seed region used for seed-based FC; the density map is overlaid on the ABA-CCFv3 template. **B)** Average seed-based FC from the dorsal striatum seed region to the rest of the brain, showing the difference between animals injected with PFF- and PBS-injected animals. **C)** Average FC between major brain parcellations (left) and circular plot of significant statistical differences (right). **Abbreviations:** ABA-CCFv3, Allen Mouse Brain Common Coordinate Framework v3; FC, functional connectivity; PBS, phosphate-buffered saline; PFF, preformed fibril; R, right hemisphere; RMS, root mean square.

To determine if the seed-level changes reflect broader regional disruptions, we then examined how correlations between major brain regions differ between groups using the ABA-CCFv3 (*28*) major structures atlas (Suppl. Fig. 1). Although the PFF-injected animals showed a global decrease in FC compared with controls (Fig 4C, right), the right dorsal striatum did not show any significant differences relative to the PBS-injected group (Fig. 4C, left). Several other structures, however, showed significant decreases in correlation both within and across hemispheres. In the PFF-injected group, the right isocortex showed significant reduction in FC with regions in the ipsilateral side such as the cortical subplate, as well as changes with bilateral regions such as the pons. The largest decrease was observed in the FC of the right thalamus with contralateral structures including the left thalamus, the hypothalamus and the left pons along with the ipsilateral cortical subplate (Fig 4C). Additionally, other correlations were also diminished, such as the correlation between the right cortical subplate and the bilateral midbrain regions and between the right medulla and the left pons.

Finally, to move beyond predefined anatomical regions, we investigated the correlation strength among major brain networks (Suppl. Fig. 8) extracted from the RABIES analysis pipeline (*29*). Five networks were assessed (Suppl. Fig. 8): the amygdala, the prefrontal cortex, the anterior somatomotor/somatosensory, the posterior somatomotor/somatosensory, and the default mode network (DMN). As opposed to the seed-based and the atlas-based analyses, the between-network analysis revealed no significant group differences (Suppl. Fig. 8).

### PS129 leads to hypoactivation during spontaneous paw movement

To examine how pS129 pathology affects motor network activity during motor activity, we used spontaneous paw movements recorded by an MR-compatible camera as a proxy for a motor task (see Methods). Although the animal’s head was fixed during acquisition, its paws moved freely (Fig. 5A). Following a spontaneous paw movement, BOLD fMRI responses across multiple structures were detected (Fig 5B left). Positive BOLD responses were observed bilaterally such as the primary and secondary motor areas, the anterior cingulate area, caudate putamen, and lateral septal nucleus of the striatum. Negative BOLD responses were observed in structures including the somatosensory regions, the retrosplenial area, and subcortically in the substantia nigra and various thalamic nuclei (Fig 5B).

**Fig 5.**
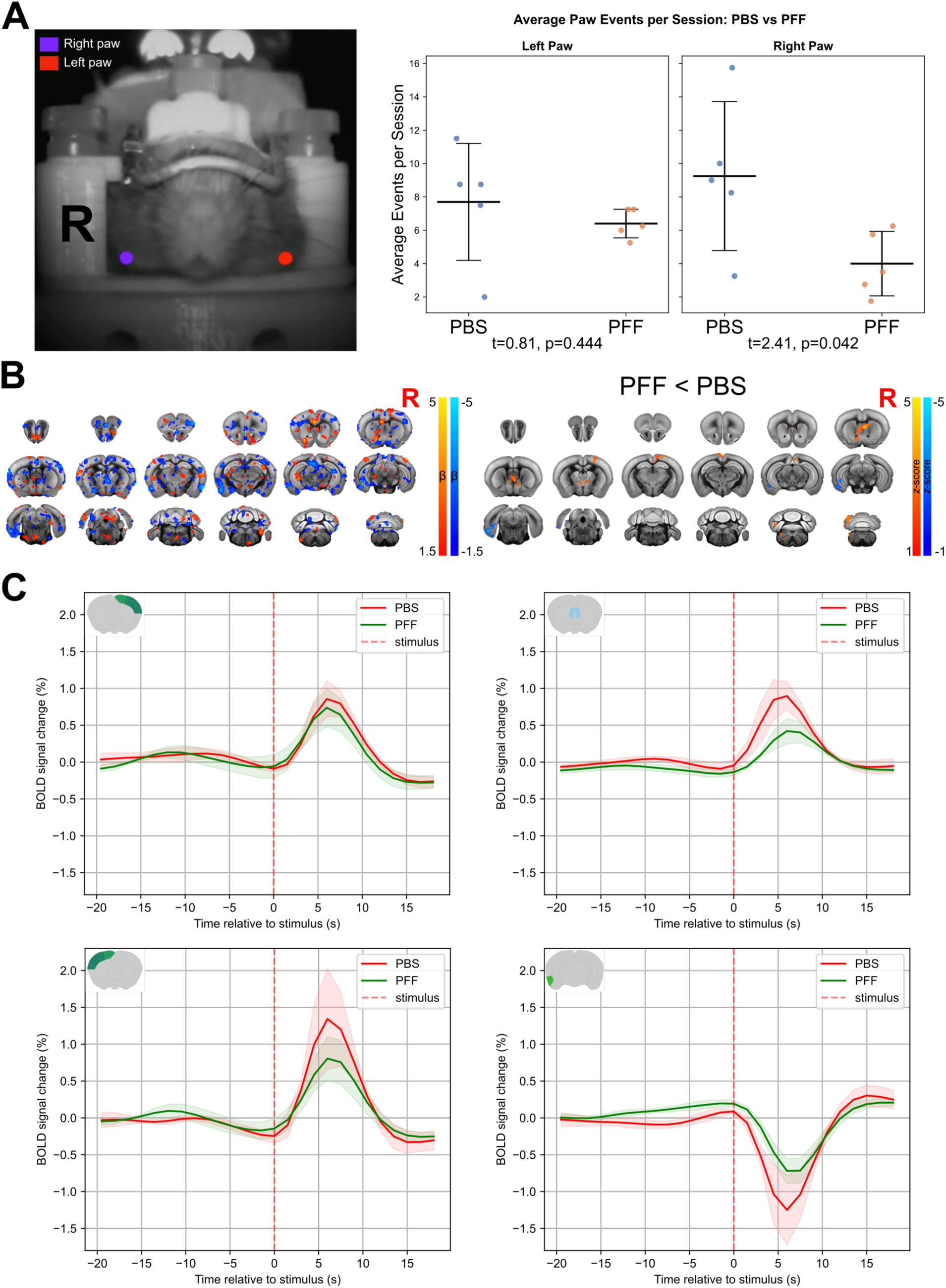
Reduced motor cortex activation in response to spontaneous left paw movements in PFF-injected animals. **A)** Anatomical landmarks used for paw tracking via DeepLabCut (left) and comparison of average movement frequency per session between groups (right). B) Group-averaged BOLD activation maps in response to left paw movements (left) and voxelwise statistical differences between groups (right). **C)** Average BOLD timeseries extracted from three regions of interest ([R MO/SS], [LSX], [L MO/SS], and [L ENT]; shading, SEM). **Abbreviations:** BOLD, blood-oxygen-level-dependent; ENT, entorhinal area; L, left hemisphere; LSX, lateral septal complex; MO, somatomotor areas; PBS, phosphate-buffered saline; PFF, preformed fibrils; R, right hemisphere; ROI, region of interest; SEM, standard error of the mean; SS, somatosensory areas.

In response to left paw (contralateral) spontaneous movement, PFF-injected animals showed reduced activation across multiple brain regions compared to controls (Fig. 5B right). In the isocortex, the somatosensory and somatomotor areas were hypoactivated in the right hemisphere, while the retrosplenial cortex showed bilateral reductions. Within the striatum, the lateral septal nucleus and caudate putamen were bilaterally hypoactivated, while the nucleus accumbens showed reduced activation only on the left side. Various motor-related nuclei in the thalamus, such as the anteromedial and ventral medial nuclei, showed bilateral hypoactivation compared to controls. On the contralateral side, regions of the pallidum and the hypothalamus showed similar reductions. Caudally, several lobules in the cerebellum showed hypoactivation in the PFF-injected group contralaterally (Fig 5B right).

In both groups, several regions of the hippocampal formation contralateral to the injection side, including the entorhinal cortex, the subiculum, the presubiculum, the dentate gyrus, and the CA1 field, exhibited negative BOLD responses. However, PFF animals showed attenuated deactivation relative to PBS controls, with smaller negative peak amplitudes (Fig. 5B).

To determine whether group differences in BOLD activation could be explained by differences in movement frequency, we compared the average number of paw movements per session between PFF-injected and PBS-injected animals (Fig. 5A right). Left paw movement frequency did not differ between groups, indicating that the group differences in BOLD activation observed during left paw movement were unlikely to be explained by differences in movement frequency. However, PFF-injected animals showed a significant decrease in right paw movement frequency compared to PBS-injected animals, yet no BOLD group differences were detected during right paw movement. The reduced movement frequency of the right paw could explain the absence of changes in response to the right paw’s movement.

Mean BOLD timeseries extracted from active voxels within anatomical regions that showed voxelwise group differences confirmed these group differences (Fig. 5C).

### PS129 covaries with different *in vivo* structural and functional measures

PS129 field fraction was correlated with structural and functional measures to assess whether structural and functional changes detected by earlier analyses covaries with pS129 pathology.

First, to test whether the pS129 accumulation follows anatomical projections from the injection site (Fig. 6A, Suppl. Fig. 9A), pS129 field fraction was correlated with the anterograde viral tracer projection volume from the AMBCA (*27*) across ipsilateral structures receiving direct projections from the dorsal striatum (Suppl. Fig. 9). This yielded a significant positive correlation *(r = 0.527, p_SA-corrected = 0.007;, 30)*, indicating that regions with stronger anatomical connectivity to the injection site tended to show greater pS129 accumulation. However, the atrophy in those regions, as presented by the W-score, showed a weak negative correlation with the field fraction (*r* = −0.124, *p*_SA-corrected = 0.538; Fig. 6D, Suppl. Fig. 9D), suggesting that stronger anatomical connection to the inoculation site does not necessarily translate to greater atrophy.

**Fig 6.**
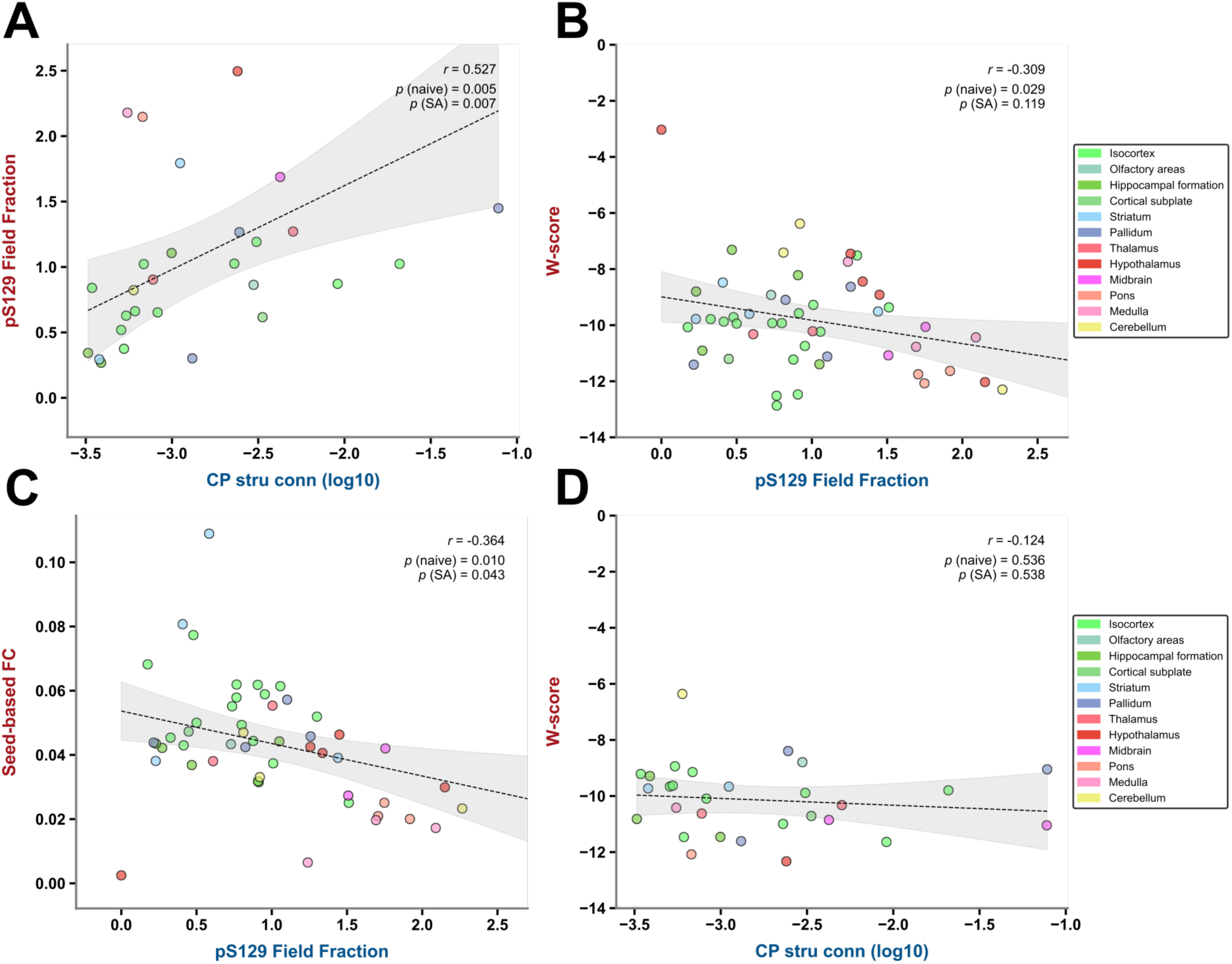
Correlations among pS129 field fraction, structural connectivity, atrophy, and functional connectivity. All panels show scatter plots with Pearson correlation coefficient (r). **A)** Structural connectivity from the dorsal striatum (CP) versus pS129 field fraction within those regions, right hemisphere. **B)** Atrophy W-scores versus pS129 field fraction, averaged across both hemispheres. **C)** Seed-based functional connectivity versus pS129 field fraction, averaged across both hemispheres. **D)** Structural connectivity from the dorsal striatum (CP) versus atrophy W-scores within those regions, right hemisphere. Dots of the same color denote substructures of a common major region; shading shows the confidence interval. Structural connectivity is derived from AMBCA anterograde tracing. **Abbreviations:** AMBCA, Allen Mouse Brain Connectivity Atlas; CP, caudoputamen; FC, functional connectivity; pS129, phosphorylated α-synuclein at S129; SA spatial autocorrelation; W-score, atrophy W-score.

Next, we examined how pS129 load correlates with the structural and functional changes reported above (see Results). For these analyses, values were averaged across hemispheres for each brain structure. Seed-based FC showed a negative correlation with the pS129 field fraction (Fig 6B, Suppl. Fig. 9) across all the brain structures (*r* = -0.364, *p*_SA-corrected = 0.043), indicating that greater pS129 burden was associated with lower FC. Similarly, W-score covaried negatively with amount of pS129 (Fig 6C, Suppl. Fig. 9) across structures (*r* = −0.309, *p*_SA-corrected = 0.119), although the corrected *p*-value did not reach the significance threshold.

## DISCUSSION

In the current body of work, we combined *in vivo* structural and functional MRI data acquired in awake, head-restrained animals with whole-brain LSFM images from the same M83 hemizygous (A53T) mice following a unilateral PFF injection in the dorsal striatum. Performing MRI in awake animals circumvented the well-documented limitations that come with anesthesia and its deleterious effects on brain networks (*31–34*). By directly correlating the pS129 pathology burden within each animal with its structural and functional MRI measures, we examined how these measures might be shaped by the underlying pathology. We also leveraged spontaneous movements during scanning to probe motor activity and how it changes between groups. To process and analyze the LSFM data, we introduced different pipelines (SPIMprep and SPIMquant) and integrated their outputs with popular MRI analysis pipelines such as RABIES (*29*). This multimodal, same-subject approach bridges a gap in synucleinopathy preclinical research where neuroimaging and histological data are acquired in separate cohorts, or the histological data does not cover the whole brain.

### Pathology spreads preferentially from the inoculation site

Mounting evidence suggests that α-synuclein pathology spreads through the brain via neuron-to-neuron transmission in a prion-like fashion (*35, 36*). An earlier staging study from Braak and colleagues (*37*) and subsequent studies demonstrating pathology spreading to grafted tissues (*38, 39*) reinforced this hypothesis in PD research. Critics have argued that this hypothesis does not explain multiple facets of the disease, including the relative sparing of nuclei synaptically connected to major disease hubs such as the substantia nigra pars compacta and locus coeruleus (*35, 40*). Instead, a cell-autonomous hypothesis has emerged to explain why some regions might be more vulnerable to pathology than others (*41*).

Inoculating M83 mice with human or mouse PFF triggers misfolding of endogenous α-synuclein and the subsequent spread of pathology to regions far from the injection site in a spatiotemporal manner (*20, 21, 23, 24, 42*). In this scenario, PFF act as a ‘template’ for new fibrils from endogenous α-synuclein that seed misfolding and aggregation. The resulting misfolded protein then propagates trans-synaptically to other regions, triggering further aggregation.

Our results show that by 120 days post-injection, pathology had spread to all major brain structures (Fig. 2, Supp. Fig. 6) as well as to the spinal cord. However, this spread was not indiscriminate. Among ipsilateral structures receiving direct projections from the injection site, pathology burden correlated positively with connection strength. Yet there was no significant difference in pathology burden between hemispheres, despite the projections from the dorsal striatum being predominantly ipsilateral.

Modelling studies using either pS129 immunoreactivity or regional MRI-derived atrophy as a proxy for pathology in humans (*43, 44*) or mice (*24, 45, 46*) have reported that a combination of cell-autonomous factors, e.g., *SNCA* or *SNCA* with *GBA* gene expression, in association with a spread mechanism represented by anatomical connectivity to the inoculation site has better predictability of pathology than either of factor alone. Our data is broadly consistent with the spread patterns reported in these studies. However, while we observed a substantial amount of pathology in regions such as the thalamus and cerebellum, others did not. Using wild-type mice rather than transgenic ones, Rahayel et al. reported that these regions are resistant to pathology (*46*). This discrepancy could be driven by the *Prnp* promoter that derives A53T expression in the M83 mice. Regions in which the *Prnp* promoter is more active would produce more A53T α-synuclein, which could also explain the pronounced pathology observed in the spinal cord and brain stem where motor neurons express *Prnp* at high levels (*47, 48*).

Other studies have argued that, while important, synaptic contacts are not the sole governing mechanism behind susceptibility to pathology. Using cell-type-specific viral tracers, Henrich et al. demonstrated that while connectivity governs the spread of pathology, its magnitude is governed by cellular phenotype rather than connection strength (*49*), with cell types such as cholinergic neurons appearing more susceptible and glutamatergic and GABAergic neurons more resilient.

These results can be interpreted in complementary ways. The absence of hemispheric differences in pathology burden, despite predominantly ipsilateral striatal projections, may reflect propagation through indirect connections over the 120-day interval. In addition, regional differences in neuronal susceptibility may influence the extent to which pathology accumulates after α-synuclein seeds reach a given region. These two explanations are not mutually exclusive. Within the ipsilateral hemisphere, however, the positive correlation between projection strength from the injection site and pS129 pathology burden supports a role for connectivity-dependent spreading, consistent with previous findings (*24, 45, 46*).

### Regional atrophy covaries with synucleinopathy, but not structural connectivity

Our DBM analysis revealed widespread neuroanatomical changes across both hemispheres in regions with different pathology burden. W-scores correlated negatively with pS129 field fraction, indicating that structures with greater pathology exhibited greater volume loss relative to controls. This correlation weakened considerably when restricted to structures receiving the strongest projections from the dorsal striatum. The dissociation between structural connectivity and atrophy suggests that atrophy is not driven by how strongly a structure is connected to the seeding epicenter but rather may reflect the selective neuronal susceptibility discussed. (*24, 43, 44*). The spatial correspondence between pS129 accumulation and atrophy is compatible with a role for synucleinopathy in neurodegeneration and cellular loss that might drive atrophy. However, the cross-sectional nature of our study precludes determining whether pathology drives atrophy, whether vulnerable regions accumulate both independently, or whether a third upstream variable contributes to both.

### PS129 inclusions disrupt regional FC, but not network-level connectivity

Rs-fMRI analysis revealed that synucleinopathy is associated with disruption in FC at the regional level. Seed-based connectivity between the injection site and the rest of the brain showed a widespread decrease in PFF-injected animals relative to controls across multiple cortical and subcortical regions. Atlas-based connectivity confirmed these findings and further demonstrated that these reductions extended beyond the seed region to include cortical-subcortical and interhemispheric connections. In primary neuronal culture, pS129 aggregates disrupt presynaptic protein machinery, impairing synaptic function and connectivity and ultimately leading to neuronal cell death (*4, 50*).

Consistent with this, Chen et al., showed that pS129 pathology selectively disrupts cortical glutamatergic transmission in the cortico-basolateral amygdala circuit (*51*). Furthermore, pS129 accumulation and the loss of axonal terminals that ensue can lead to a reduced synaptic transmission even before synaptic loss or cell death (*52*). Our whole-brain analysis extends these observations, showing that regions bearing greater pathology burden show weaker functional connectivity to the injection site. However, these previous *ex vivo* findings were obtained in wild type C57BL/6J mice, and their applicability to the M83 transgenic model used here should be interpreted with caution. By contrast, FC between large-scale networks derived from the ICA analysis, such as the DMN and the somatosensory/somatomotor networks, did not yield significant group differences. This dissociation may reflect several factors: the effects of pS129 pathology on FC may operate at the regional rather than the network level. Alternatively, it may indicate that not all connections between circuits are affected equally (*51*), or that the coarseness of the parcellations is too high to detect changes within the subnetworks. Indeed, Dipasquale and colleagues reported that, in Alzheimer’s disease, high-dimensional ICA decomposition revealed within-network FC alterations that were absent in low-dimensional analysis (*53*).

### α-synuclein inclusions lead to hypoactivation in a cell-autonomous manner

We leveraged the spontaneous paw movements in our awake animals during image acquisition as a proxy for a motor task. Our results showed hypoactivation in somatomotor/somatosensory regions, as well as in the medial septum and cerebellar cortex in response to PFF seeding. The dissociation between left paw movement (no frequency difference between groups, yet BOLD hypoactivation in PFF animals) and right paw movement (reduced frequency, no BOLD difference) suggests that abnormal neuronal activity is more likely to explain these results than movement frequency differences. The reduced number of right paw movement events in PFF animals (∼x2 reduction) may also yield limited statistical power to detect ipsilateral BOLD differences.

Using a variety of motor tasks such as button presses or finger tapping, multiple studies have characterized motor activation in PD patients using BOLD fMRI or positron emission tomography (PET). Despite conflicting results, a recent meta-analysis showed a pattern of consistent hypoactivation in the posterior putamen, the primary motor cortex, the supplementary motor area, and the cerebellum (*54*). Our findings extend these observations to a preclinical model. However, it is worth noting that those tasks were voluntary and followed a predefined paradigm, while our results come from spontaneous movements that did not involve a cue. Various studies, both *in vivo* and *ex vivo*, have attempted to unravel the mechanism behind this hypoactivation. The classical rate model attributes motor dysfunction in PD to a reduction in excitatory output from the thalamus to the sensorimotor cortex, resulting from excessive inhibitory output of the globus pallidus internus following nigrostriatal dopamine loss (*55, 56*). This model ascribes cortical hypoactivation to suppression of thalamic output while the cortex itself remains intact. Our data, however, suggests that another mechanism may be at play. We observed substantial α-synuclein inclusions in the motor cortex and neighboring areas, indicating that the motor cortex is directly affected by synucleinopathy. Consistent with this, Chen et al. demonstrated that pS129 aggregates lead to hyperexcitability in pyramidal cortical neurons through a cell-autonomous mechanism, accompanied by increased input resistance and decreased cell capacitance, leading to a disrupted excitation/inhibition ratio (*57*). Moreover, those aggregate-bearing neurons showed shorter dendrites, loss of dendritic spines, and reduced glutamatergic output (*57*). A recent study showed that pS129 aggregates in M2 presynaptic glutamatergic neurons are sufficient to impair glutamatergic transmission onto striatal spiny projection neurons, independent of cortical neurons or striatal dopamine terminals degeneration (*58*).

The BOLD response predominantly reflects synaptic input and intracortical processing rather than spiking output of individual neurons (*59*). Accordingly, a cortex with hyperexcitable but synaptically compromised neurons would be expected to generate a weaker hemodynamic response. A third possible explanation is spinal cord involvement, since pS129 pathology was present in the spinal cord of PFF-injected animals. However, the cell-autonomous mechanism offers a more parsimonious account, as it also explains the observed decreases in FC in cortico-thalamic and cortico-amygdala networks (*51, 52, 57, 60*).

The differences in the negative BOLD response between groups observed in areas within the hippocampal formation of PFF-injected animals may similarly reflect disrupted inhibitory GABAergic transmission. Negative BOLD responses have been linked to decreased neuronal activity *(61, see 62 for a review)* and to increased GABA concentration (*63*). This interpretation, however, warrants further investigation.

Motor cortex hypoactivation, if clinically validated, could serve as a non-invasive imaging-based biomarker for preclinical and prodromal PD and other types of synucleinopathy. Future studies should assess its utility in terms of differential diagnosis, correlation with disease progression and severity, and prediction of therapeutic outcome.

### PS129 inclusions are involved in structural and functional changes

The identification of α-synuclein as the main component of LBs and LNs (*1*) and the revelation that ∼90% of α-synuclein within these inclusions is phosphorylated at serine 129 (*2, 3*) sparked a longstanding debate over whether pS129 and these insoluble aggregates play a toxic or protective role. Some studies argued that pS129 and its aggregates are not inherently toxic and may even exert a neuroprotective effect by preventing further aggregation or sequestering toxic α-synuclein species (*8, 9, 64*), while others showed that pS129 is toxic and contributes to neuronal cell death (*11*). Our findings show that greater levels of pS129 inclusions are associated with both regional volume loss and decreased functional connectivity, a pattern consistent with a deleterious role for pS129 aggregates, or components of the inclusion that accumulate with aggregates. However, the cross-sectional nature of the study does not permit determination of whether pS129 accumulation preceded or followed structural and functional changes.

Several interpretations merit consideration. Phosphorylation at S129 may follow rather than drive aggregation (*9*), and the mature inclusions themselves may serve a protective role by sequestering more toxic, soluble α-synuclein species (*64*). Clinically, LB density does not consistently correlate with disease severity, cognitive symptoms, or disease duration in synucleinopathies (*65*), further supporting the argument that inclusions are not the principal drivers of neurodegeneration. Consistent with this view, Mahul-Mellier et al. demonstrated that it is the process of inclusions formation, through sequestering protein transport machinery, rather than the mature inclusions themselves, that drives cellular toxicity (*66*). Furthermore, several groups have demonstrated that soluble prefibrillar species, including oligomers and protofibrils are the primary toxic agents (*67–70*).

While our data cannot distinguish among these possibilities, the spatial correspondence between pS129 burden and the structural and functional changes suggests that regions bearing higher pathology load are more vulnerable to neural injury, regardless of whether pS129 inclusions drive toxicity alone or in conjunction with other α-synuclein species.

### Conclusions

In conclusion, using a same-subject multimodal dataset combining whole-brain MRI and LSFM, our study demonstrates how pS129 pathology correlates with mesoscale structural and functional changes. Following a unilateral PFF injection in the dorsal striatum, pathology was detected throughout the brain and spinal cord, with preferential deposition in the midbrain. This pathology spread was governed by anatomical connectivity to the seeding location and was correlated with diminished functional connectivity, regional atrophy, and hypoactivation in task-evoked BOLD response. The correspondence between pS129 accumulation and structural and functional degredation are consistent with a toxic role for pS129 inclusions. Our study provides a framework for understanding the correlation between cellular and mesoscale changes and lays the groundwork for evaluating therapeutics.

### Limitations and future directions

While our study presents a unique dataset combining whole-brain LSFM with awake functional and structural MRI within the same animals, several limitations should be acknowledged. Although, we have demonstrated that correlations are consistent within animals, the small sample size, constrained by the logistical challenges of acquiring LSFM and awake MRI data from the same animals, limited our ability to detect changes with smaller effect sizes. We opted to use males only because males show more aggressive neurodegeneration (*21*) and males are more prone to develop PD (*71*). The AMBCA structural connectivity data (*27*) used to test pS129 spreading are derived from anterograde viral tracing vectors, leaving the retrograde and bidirectional connectivity uncharacterized. Further studies are warranted to test the trans-synaptic vs other extracellular mechanisms. Finally, all measures are cross-sectional and derived from a single-time point. A longitudinal study would be necessary to determine the temporal relationship between the pS129 accumulation and the development of structural and functional changes.

## Materials and Methods

### Study Desing

In the current study, we combined *in vivo* MRI with whole-brain LSFM to investigate structural and functional changes in M83 hemizygous male mice following either a seeding injection of human preformed fibrils (hu-PFF) of α-synuclein or phosphate-buffered saline (PBS) in the dorsal striatum. To mitigate the confounding effects that anesthesia has on functional networks (*31–34*), all MR imaging was performed in awake, head-fixed animals. Furthermore, we examined the relationship between pS129 pathology, quantified by LSFM, and mesoscopic changes observed using MRI in order to test whether pS129 pathology underlies these alterations.

### Ethics

All animal experiments were performed in compliance with the ARRIVE guidelines.

### Animals

10 transgenic hemizygous M83 male mice expressing one copy of human α-synuclein with the A53T familial mutation under the control of the mouse prion protein promoter were used.

### PFF and PBS injection

Intracerebral injection of α-synuclein PFF was performed to accelerate toxic accumulation of α-synuclein aggregates that precipitate earlier disease onset (*4, 72*). Animals between 4 and 6 months old were injected with either recombinant hu-PFF (5 animals) *(for more details about PFF preparation and characterization, see 73, 74)* or PBS to serve as control (Fig. 1A). Surgeries were performed as previously described in (*23*). Briefly, the animals were anesthetized using isoflurane and then positioned on a stereotaxic platform. Each mouse received a single injection of either 2.5 μl of PFF (12.5 μg from 5 mg/ml PFF) or 2.5 μl of sterile PBS into the dorsal striatum of the right hemisphere.

### Head-post surgery

See supplementary method.

### MRI acquisition

Scanning sessions were performed on a 9.4 T small animal scanner. T2*-weighted images for fMRI were acquired using a gradient-echo EPI (GE-EPI) sequence covering the whole brain with the following parameters: TE/TR = 12/1500 ms, flip angle (FA) of 60°, FOV of 19.2 × 9.6 × 15 mm, resolution 300 × 300 × 500 µm^3^. For anatomical images, a three-dimensional magnetization transfer gradient echo sequence (MT-GRE3D; FLASH) was used. Acquisition parameters were as follows: TE/TR = 3.54/40 ms, FA = 9°, isotropic voxel size = 150 µm, FOV = 19.2 × 9.6 × 19.2 mm. Animals were continuously during scanning using an MR-compatible camera. Video footages were recorded and synchronized with the functional scans acquisition.

### Tissue collection and clearing

Following MRI, seven animals (5 PFF, 2 PBS) were perfused transcardially with heparinized PBS and 4% PFA. After extraction, brains were preserved using SHIELD (LifeCanvas Technologies; LCT; Cambridge, MA, USA) (*75*), delipidated using LCT Clear+ Delipidation Buffer, and immunolabeled per SmartBatch+ protocol (v5.06; LCT) with rabbit anti-human α-synuclein (EP1536Y; Abcam) detected by donkey anti-rabbit SeTau-647 (LCT), alongside propidium iodide as a nuclear counterstain. Samples were refractive index-matched (EasyIndex, RI = 1.52; LCT) and imaged on an UltraMicroscope Blaze LSFM (Miltenyi Biotec) at 2.5 µm z-step and 1.625 × 1.625 µm x-y resolution (561 nm/647 nm excitation).

### LSFM data processing

Raw stitched images in imaris format (ims) were converted to OME-ZARR (ome.zarr) format using SPIMprep (v0.1.0-alpha) (https://github.com/khanlab/SPIMprep). SPIMquant (v0.5.0-alpha1) (https://github.com/khanlab/SPIMquant) was used for downstream processing. Whole-brain masks for the PI channel were generated using k-class Gaussian Mixture Model (GMM) segmentation. Following ANTs N4BiasFieldCorrection (*76*), the PI channel was registered to the anatomical image of the Allen Mouse Brain Common Coordinate Framework v3 atlas *(ABA-CCFv3; 28)* using linear and nonlinear registration. The pS129 channel was bias-field corrected and segmented by intensity thresholding. Field fraction (percentage of positive voxels per region) was computed at two parcellation granularities (12 and 51 ROIs per hemisphere; Suppl. Figs. 1–2). Hemispheres were pooled after confirming no significant laterality differences (Suppl. Fig. 3).

### Spinal cord samples preparation and analysis

See supplementary methods.

### Deformation-based morphometry (DBM)

The anatomical images were denoised, bias-field corrected, and brain-maksed. The processed images were used to construct an unbiased study-based template through successive rounds of linear and nonlinear registration with continuous template refinement using ANTs registration algorithm (*77*). DBM analysis was performed using optimized_antsMultivariateTemplateConstruction pipeline *(78;* https://github.com/CoBrALab/optimized_antsMultivariateTemplateConstruction/). The relative Jacobian images were smoothed with a 4-voxel (0.6 mm) full width at half maximum (FWHM) Gaussian kernel and used for voxelwise statistical analysis. The mean and standard deviation of an age-matched control population (*24*) were used to calculate W-score images (*79*) from the full Jacobian images. Mean W-score values within each parcel of the fine-grained atlas (see above) were calculated and used to investigate how DBM-derived volume changes vary with other measures.

### Functional MRI preprocessing and confounds correction

RABIES pipeline was used for fMRI images preprocessing and confounds correction *(v0.5.0; 29)*. The anatomical images were used to construct an unbiased, study-based template The unbiased template was then registered to the DSURQE ex-vivo MRI atlas (*80–82*). The functional images were motion-corrected, distortion corr3ected using corresponding anatomical images, and were finally resampled to the reference atlas common space at a voxel resolution of 0.2 × 0.2 × 0.2 mm^3^. The timeseries were then detrended and band-pass filtered using a cutoff of 0.01 Hz. The six rigid body motion parameters estimated earlier were regressed out of the data. Finally, timeseries were converted to percent blood-oxygen-level-dependent (BOLD) signal fluctuations by dividing by the grand mean signal across voxels and multiplying by 100.

To allow use of the Allen Mouse Brain Atlas (ABA), the anatomical image of the ABA-CCFv3 was registered to the DSURQE atlas (Suppl. Fig. 4). The coarse (Suppl. Fig. 1) and fine parcellations (Suppl. Fig. 2) were resampled to the space and voxel resolution of the functional images.

### Structural connectivity

See supplementary methods.

### Seed-based FC

Cleaned timeseries were smoothed using a FWHM Gaussian kernel of 0.5 mm. The seed region was defined from the structural-connectivity density map to cover the approximate injection area (Fig. 4A, Suppl. Fig. 5). Pearson correlations between seed and whole-brain voxel timeseries were Fisher *z*-transformed for voxelwise analysis. Group-average FC maps were generated per group, and mean connectivity within each fine-grained atlas parcel was extracted for subsequent correlation analyses.

### Atlas-based functional connectivity (FC)

Pairwise Pearson correlations between coarse atlas parcels, calculated from cleaned unsmoothed time series and *z*-transformed, were averaged across runs to produce subject-level connectivity matrices for group-level statistical inference.

### Independent component analysis (ICA)

The smoothed data from all subjects were concatenated and decomposed into 30 networks using RABIES’ precomputed networks as priors. Between-network Pearson correlations among five major networks (Suppl. Fig. 8) were *z*-transformed and used for group analysis.

### Paw movement tracking

Bilateral paw positions (Fig. 5A) were tracked across all runs using DeepLabCut (*83*). Per-paw coordinates were mean-centered, and movement events were detected as peaks in the first principal component of paw motion. Event timing was used to construct per-paw regressors convolved with a double-gamma hemodynamic response function (HRF) and temporal derivative for voxelwise GLM analysis. Group-level inference employed mixed effects (flameo) with cluster-forming threshold Z > 2.1 and cluster-level correction at p < .05. Regional BOLD time series were extracted from atlas-defined masks (Fig. 5C), and stimulus-locked responses were averaged within ±20 s windows around movement events across runs and subjects. Per-session movement frequency was compared between groups.

### Correlation analysis

See supplementary methods.

### Statistical analysis

For voxelwise DBM and seed-based FC analyses, group differences were assessed using two-sample tests with non-parametric permutation testing (FSL PALM; (*84*); 10,000 permutations) and threshold-free cluster enhancement (TFCE; p < .05). Atlas-based and ICA-based FC matrices were compared using unpaired two-sample permutation tests. Hemispheric pS129 field fraction differences were tested with paired t-tests (FDR-corrected), and paw movement frequency with unpaired t-tests. Subject-level correlations were evaluated via one-sample t-tests on Fisher z-transformed coefficients. Group-level spatial correlations were assessed against spatially autocorrelation-preserving surrogate maps (*30*).

## Acknowledgements

Funding

## List of Supplementary Materials

Materials and Methods

Fig. S1 to S9

Movie S1 to S2

**Supplementary Video 1.** Whole-brain pS129 immunostaining in PBS- and PFF-injected M83 mice. Three-dimensional rendering of LSFM volumes showing pS129 (yellow) and PI (blue) in a PBS-injected control (left) and a PFF-injected mouse (right). The PFF-injected brain displays widespread pS129 pathology throughout cortical and subcortical regions, whereas the PBS-injected brain shows minimal signal. **Abbreviations:** LSFM, light-sheet fluorescence microscopy; PBS, phosphate-buffered saline; PFF, preformed fibrils; PI, propidium iodide; pS129, phosphorylated-s129-α-synuclein.

**Supplementary Video 2.** Brain and spinal cord pS129 pathology in a PFF-injected M83 mouse. Three-dimensional rendering of the intact brain and spinal cord LSFM from a single PFF-injected subject, showing the spatial distribution of pS129 α-synuclein immunostaining throughout the CNS. **Abbreviations:** CNS, central nervous system; LSFM, light-sheet fluorescence microscopy; PBS, phosphate-buffered saline; PFF, preformed fibrils; PI, propidium iodide; pS129, phosphorylated-s129-α-synuclein.

## Author contributions

Conceptualization: RSM, MAMP, ARK, AE

Methodology: AE, VN, SB, KS, ARK, VFP, ST, MMC, WL, IS, TMD

Investigation: RSM, MAMP, ARK, AE

Visualization: AE, ARK, SB, KS

Funding acquisition: RSM, MAMP, ARK

Supervision: RSM, MAMP, ARK, VFP

Writing – original draft: AE

Writing – review & editing: AE, RSM, MAMP, ARK, VFP, SB, KS

## Data and materials availability

All the MRI data used in this manuscript is publicly available (https://osf.io/8adfe), the LSFM data is available in physical format upon request. The code used for analysis is available here: https://github.com/amrka/MRI-LSFM-Synuclein-manuscript.

## Materials and Methods

### Ethics

All animals were bred and housed at Western University’s animal facility. All experiments were conducted according to the Canadian Council on Animal Care (CCAC) guidelines and were approved by the Animal Care and Veterinary Services (ACVS) at the Western University (protocols: #2020-162, #2020-163, and #2021-160). All experiments were performed in compliance with the ARRIVE guidelines.

### Animals

Ten transgenic hemizygous M83 male mice expressing one copy of human α-synuclein with the A53T familial mutation under the control of the mouse prion protein promoter (B6; C3-Tg(Prnp-SNCA*A53T) 83Vle/J, RRID: IMSR_JAX:004,479) on the C57/C3H background were used (*23, 47*). The animals were bred in house and maintained on the C57BL/C3H background. Mice were housed in standard plexiglass cages with ad libitum access to Harlan food and water. Containing rooms were controlled for temperature and humidity (22-25°C and 40-60%) and were operated on light/dark cycle from 7am to 7pm. Animals were group-housed (2-4 animals per cage). Mice were housed individually whenever aggressive behavior was observed. All control mice used were littermates. Epidemiologically, PD is twice as common in males than in females (*85*) and in this study male mice were used exclusively as it has been demonstrated that they exhibit a more aggressive phenotype compared to female counterparts (*21*).

### PFF and PBS injections

It has been determined that M83 hemizygous mice (TgM83+/-) do not present PD phenotypes until old age (22-24 months) (*47*), therefore intracerebral injection of α-synuclein PFF was performed to accelerate toxic accumulation of α-synuclein aggregates that precipitate earlier disease onset (*4, 72*).

Animals between 4 and 6 months old were injected with either recombinant hu-PFF (5 animals) or PBS to serve as control (Fig. 1A). Dynamic light scattering (DLS) was used to characterize fibrils to ascertain the average diameter of PFF was <100 nm *(73, for more details about PFF preparation and characterization, see 74)*. Mice were anaesthetized with isoflurane (4-5% for induction, 1.8-2% for maintenance) delivered in O_2_ at flow rate of 0.8 L/min. Prior to incision, mice received Metacam (Meloxicam; 5 mg/kg; NADA 141-219) and Bupivacaine (Bupivacaine hydrochloride; 5 mg/kg; NADA 018053) subcutaneously at the incision site for analgesia. Mice were then positioned on a stereotaxic platform, and a small incision was made in the scalp to expose the skull and the midline. Each mouse received a single injection of either 2.5 μl of PFF (12.5 μg from 5 mg/ml PFF) or 2.5 μl of sterile PBS into the dorsal striatum of the right hemisphere at the following coordinates relative to bregma: +0.2 mm anteroposterior (AP), +2.0 mm mediolateral (ML), and -2.6 mm dorsoventral (DV). A Hamilton syringe was initially lowered to a depth of 3.0 mm and then retracted to -2.6 mm before injection. The flow rate of the injected material was maintained at 250 nl/min. The syringe was left in place for 1-2 minutes after finishing the injection and then slowly withdrawn. The incision was then sutured, and mice received 1 ml injection of 0.9% saline subcutaneously before being allowed to recover under a heat lamp. Once recovered, mice were returned to their cage and monitored closely for two weeks. Mice received a daily injection of Metacam (5 mg/kg) intraperitoneally for two days after the surgery. No significant health decline or side effects were noted in any animal following the surgery.

### Head-post surgery

To enable MRI-awake imaging, a head-post-implanting surgery was performed 15-17 weeks after PFF or PBS injection. The surgery and acclimation were performed as previously described (*86*). Briefly, the anesthesia was initiated as previously described (see above), a small incision was made in the scalp, and a polycarbonate MRI compatible 3D printed head-post was fixed to the skull using dental cement and cured using UV light. Following a recovery period, the animals were acclimated to the scanner environment over 9 days using a mock scanner with increasing duration of restraining and noise level. Following the head-post implantation surgery, the animals were individually housed to avoid damage to the head-posts.

### MRI acquisition

Once the animals were acclimated to the scanner environment, MRI scanning was conducted. One scanning session was acquired from each animal containing anatomical and functional scans. Scanning sessions were performed on a 9.4 T small animal scanner with a 31 cm bore magnet equipped with a 6 cm Magnex HD gradient coil insert of 1 T/m strength (Agilent, Palo Alto, CA, USA) interfaced to a Bruker Avance Neo console with software package of Paravision-360.3.3 (Bruker BioSpin Corp, Billerica, MA), and a single-loop surface coil of 2 × 1 cm^2^. T2*-weighted images for fMRI were acquired using a gradient-echo EPI (GE-EPI) sequence covering the whole brain with the following parameters: TE/TR = 12/1500 ms, flip angle (FA) of 60°, FOV of 19.2 × 9.6 × 15 mm, in-plane resolution 300 × 300 µm^2^, and slice thickness of 500 µm. Four runs of 400 volumes were acquired consecutively for a total scan time of 40 min. For anatomical images, a three-dimensional magnetization transfer gradient echo sequence (MT-GRE3D; FLASH) was used. Acquisition parameters were as follows: TE/TR = 3.54/40 ms, FA = 9°, isotropic voxel size = 150 µm, FOV = 19.2 × 9.6 × 19.2 mm, number of averages = 4. Total acquisition time was approximately 26 minutes.

The animals were continuously monitored throughout scanning using a 12M-i newSensor MR-compatible camera equipped with an infrared (IR) LED (MRC Systems GmbH, Heidelberg, Germany). Video footages were recorded during fMRI acquisition using Python OpenCV library (https://opencv.org/) running on a Raspberry Pi 4 Model B (Raspberry Pi Ltd., Cambridge, UK) at a frame rate of 20 frames per second (fps). Camera recording was synchronized to the onset of each functional run via a transistor-transistor logic (TTL) pulse triggered by the scanner.

### Tissue collection and clearing

Following scanning, 7 animals (5 injected with PFF and 2 injected with PBS) were anesthetized using a mixture of ketamine (100 mg/kg) and xylazine (20 mg/kg) in 0.9% sterile saline. Animals were then transcardially perfused with ice-cold heparinized 1X PBS followed by cold 4% PFA. The brains were then carefully extracted from the skulls, washed with 1X PBS, and stored in 1X PBS containing 0.02% sodium azide at 4°C for 24 hours on an orbital shaker. Sample preservation was done via intramolecular epoxide linkages to prevent degradation using SHIELD reagents according to the manufacturer’s instructions (LifeCanvas Technologies; LCT; Cambridge, MA, USA) (*75*). Delipidation was performed using LCT Clear+ Delipidation Buffer. Whole brain samples were first incubated in the buffer at 45°C for 24 hours, followed by passive delipidation for a total delipidation time of 9 days. Following delipidation, the samples were immunolabelled according to LCT Active Labelling SmartBatch+ protocol (v5.06; LCT). Samples were incubated with 72 µg (∼10 μg per brain) of rabbit anti-human α-synuclein (EP1536Y; Cat#ab51253, lot# 1055686-1; Abcam), 288 µl (∼41 μg per brain) of Propidium Iodide (PI) (Cat#P3566; Invitrogen), and anti-human α-synuclein was detected using a donkey anti-rabbit IgG SeTau-647 (∼20 μg per brain) secondary antibody (Cat#DkxRb-ST; LCT). Samples were subsequently refractive index-matched by sequential incubation in 50% EasyIndex (RI = 1.52; LCT) for 24 hours at 37°C, followed by 100% EasyIndex (RI = 1.52; LCT) for 24 hours at 37°C. Whole-brain volumetric imaging was performed on an UltraMicroscope Blaze LSFM (Miltenyi Biotec) using a 4x/0.068 NA objective with a 2.5 µm z-step and 1.625 × 1.625 µm x-y pixel size. PI and pS129 signals were acquired at 561 nm and 647 nm excitation wavelengths.

### LSFM data processing

Raw tiled images stack in OME-TIFF (ome.tif) format for the PI and α-synuclein were stitched offline using Imaris (Bitplane, Oxford Instruments) and exported in imaris format (ims).

Two in-house Snakemake-based workflows were used to process the images. First, SPIMprep (v0.1.0-alpha) (https://github.com/khanlab/SPIMprep) was used to convert stitched data to OME-ZARR (ome.zarr) format and to organize the data according to Brain Imaging Data Structure (BIDS) Microscopy specification (*87*).

SPIMquant (v0.5.0-alpha1) (https://github.com/khanlab/SPIMquant) was used for processing, which included the following steps. OME-ZARR images were imported and converted to Nifti format using ZarrNii v0.17.1-alpha1 (https://github.com/khanlab/zarrnii), using multi-resolution level 3 (8x downsampled) for downstream registration and level 0 for anatomical segmentation steps. Whole-brain masks for the PI channel were generated using k-class Gaussian Mixture Model (GMM) segmentation implemented in ANTs Atropos (*88*), with k=9, assigning all classes having at least 50% volumetric overlap with an affine-warped spatial prior to the brain mask foreground class. Following ANTs N4BiasFieldCorrection(*76*) the PI channel was registered to the anatomical image of the Allen Mouse Brain Common Coordinate Framework v3 atlas *(ABA-CCFv3; 28)*. The registration was done in two stages; linear registration using 12 degrees of freedom (dof), followed by nonlinear deformable registration. Both stages were performed using greedy registration software (https://github.com/pyushkevich/greedy). The inverse transformations from both stages were used to warp the parcellation atlases from ABA-CCFv3 into subject space for region-based analysis.

In preparation for α-synuclein field fraction quantification, N4 bias field correction was applied to the α-synuclein channel using ANTs N4BiasFieldCorrection (*76*). Bias field maps were first estimated at downsampling level 3 (8× isotropic downsampling relative to full resolution), then the correction was applied at full resolution by upsampling the estimated bias field and dividing it out of the original data. This step aims to eliminate smoothly varying intensity changes across the entire brain, so that a single global threshold can be effectively applied on the corrected image.

Segmentation of α-synuclein was performed using intensity thresholding with fixed threshold value; voxels exceeding the threshold value were considered positive. Atlas parcellations warped into subject space were used to extract per-region quantitative metrics. Two parcellations with different levels of granularity were used: a coarse one comprising 12 regions of interest (ROIs) per hemisphere (Suppl. Fig. 1) and a fine one comprising 51 ROIs per hemisphere (Suppl. Fig. 2). For statistical analyses, field fraction was used as the primary metric. Field fraction is defined as the percentage of positive voxels within each brain region. PS129 field fraction between hemispheres did not show any statistically significant differences (Suppl. Fig. 3), so we pooled both hemispheres together for all subsequent analyses. Quality control (QC) reports generated by SPIMquant were visually inspected for all subjects to verify the accuracy of each processing step, including registration, segmentation, and quantification.

### Spinal cord samples

To explore the spread of pS129 to the spinal cord, six M83 male mice age (5-6 months) were injected with α-synuclein PFF as previously described (see above). 3-4 months after injection, the 6 animals, in addition to two age-matched control animals, were perfused and the brain and the spinal cord were carefully extracted and preserved as described above.

### Spinal cord clearing

The brain with the spinal cord samples were processed using the same SHIELD preservation and refractive index-matching protocol described for the brain samples (see Main Methods), with the following modifications. Delipidation was performed using LCT Clear+ Delipidation Buffer by passive incubation for 2.5 days, followed by active delipidation in SmartBatch+ buffer for 30 hours. Six spinal cord samples were immunolabelled as a single batch using reagent volumes scaled for 12 samples according to the LCT Active Labelling SmartBatch+ protocol (v5.06; LCT): 72 µg of rabbit anti-Alpha-Synuclein (phospho S129) primary antibody (EP1536Y; Cat#ab51253, lot# 1055686-1; Abcam), 288 µl of Propidium Iodide (Cat#P3566; Invitrogen), and 144 µg of donkey anti-rabbit IgG SeTau-647 secondary antibody (Cat#DkxRb-ST; LCT). Following refractive index matching, samples were washed in PBS containing 0.1% sodium azide (PBS-N) for 48 hours with two buffer exchanges prior to imaging. Volumetric imaging was performed on an UltraMicroscope Blaze LSFM (Miltenyi Biotec) using a 1x/0.034 NA objective with a 100 µm z-step and 5.9 × 5.9 µm x-y pixel size. PI and pS129-α-synuclein signals were acquired at 561 nm and 647 nm excitation wavelengths, respectively. Only spinal cord data were used for subsequent image analysis in the present study.

### Spinal cord data analysis

The PI and α-synuclein pS129 images were processed similarly using SPIMprep and SPIMquant (see above). To correct for the curvature of the spinal cord during acquisition, spinal cord straightening was performed in two sequential passes (Fig. 2D). In the first pass, the masked PI volume was split into 2D slices, and the center of gravity (CoG) of each slice was calculated. A CoG point was retained only if it was located beyond a minimum Euclidean distance from the mask boundary. The CoG coordinates were saved as a sparse NIfTI label image and used as the centerline input for the Spinal Cord Toolbox *(SCT; 89)*. SCT’s sct_straighten_spinalcord was used to produce an initial straightened volume. This intermediate volume was further split into 2D slices and the CoG coordinates were similarly calculated and filtered before getting passed into a second round of sct_straighten_spinalcord to produce a final straightened spinal cord PI volume. The second round was warranted in most cases to correct for residual curvature remaining after the first pass. A representative straightened volume was used as template and was manually segmented into cervical, thoracic, lumbar, and sacral/coccygeal regions (Fig. 2D). All subjects were length normalized to this template using ANTs (*77*). The two transformations used to straighten the PI volume, in addition to the template length normalization, were concatenated and applied to the field fraction output image from SPIMquant, and the field fraction within each region was quantified.

### Deformation-based morphometry (DBM)

For the DBM analysis the MT-GRE3D anatomical images were denoised and bias-field corrected iteratively using adaptive non-local means (ANLM) algorithm (*90*) and N4 (*76*) prior to registration. A brain mask was then generated by affine registration of each image to the DSURQE atlas (*80–82*) and used to inform a final round of weighted bias field correction before DBM analysis.

The preprocessed images were used to construct an unbiased study-based template through successive rounds of linear and nonlinear registration with continuous template refinement using ANTs registration algorithm (*77*). The DSURQE atlas was used as the final target to anchor the resulting template in a common anatomical reference space.

DBM analysis was performed using optimized_antsMultivariateTemplateConstruction pipeline *(78;* https://github.com/CoBrALab/optimized_antsMultivariateTemplateConstruction/). The anatomical image from each subject was linearly and nonlinearly registered to the group template. Log-Jacobian determinant images were computed from the nonlinear warp field using the geometric estimator, such that positive values indicate local volume expansion and negative values denote contraction relative to the group template. Relative Jacobians were derived by removing the bulk affine component embedded within the nonlinear warp field, isolating local shape differences independent of global brain size. Full Jacobians, reflecting total volumetric differences including global scaling, were computed by combining the affine and nonlinear Jacobians. The relative images were smoothed with a 4-voxel (0.6 mm) full width at half maximum (FWHM) Gaussian kernel to meet assumptions for statistical testing.

The average and standard deviation images of age-matched *ex-vivo* control population injected with PBS (*24*) were used to normalize the unsmoothed full Jacobian maps. A W-score which quantify regional volume deviation from the control population was calculated (*79*). The mean image of the control population was subtracted from the full Jacobian map of each PFF-injected animal and divided by the standard deviation image of the control population to generate a normalized map. Mean W-score values within each parcel of the fine-grained atlas (see above) were calculated and used to investigate how DBM-derived volume changes vary with other measures.

### Functional MRI preprocessing

The open-source software RABIES pipeline was used for fMRI images preprocessing *(v0.5.0; 29)*. After an initial motion realignment step, a trimmed average across all EPI frames was generated. Each EPI frame was rigidly aligned to this mean image to estimate head motion parameters. Anatomical images from all sessions and subjects were used to construct an unbiased, study-based template through iterative nonlinear registration to the dataset consensus average (*77*). The final form of the unbiased template was registered to the DSURQE ex-vivo MRI atlas (*80–82*) using nonlinear registration. For EPI susceptibility distortion correction, each EPI volume was nonlinearly co-registered to its corresponding anatomical image to recover the underlying brain geometry (*91*). To resample the EPI volumes to the common space, all transformations, including, head motion correction, susceptibility distortion correction, and common space alignment, were concatenated and applied in a single shot to avoid multiple interpolations (*92*). The preprocessed timeseries were finally resampled to the reference atlas common space at a voxel resolution of 0.2 × 0.2 × 0.2 mm^3^.

### Confound correction

Confound correction was done using RABIES (*29*). Preprocessed timeseries were first detrended to remove first-order drifts and mean signal. A third-order Butterworth high-pass filter with a cutoff frequency of 0.01 Hz was then applied to remove slow scanner drifts and other low-frequency non-neural fluctuations. Nuisance regressors were high-pass filtered using the same filter prior to regression to preserve orthogonality between temporal filtering and confound regression and to avoid the reintroduction of the nuisance signal previously removed from the data (*93*). The six rigid-body head motion parameters estimated earlier were modelled at each voxel using ordinary least squares and regressed out of the data. Finally, timeseries were converted to percent blood-oxygen-level-dependent (BOLD) signal fluctuations by dividing by the grand mean signal across voxels and multiplying by 100.

### Structural connectivity

To examine how the structural projections from the disease epicenter (the injection site) correlate with the spread of the α-synuclein PFF and the volumetric changes that follow, we leveraged the anterograde viral tracing data from Allen Mouse Brain Connectivity Atlas *(AMBCA; 27)* as a proxy for structural connectivity. Injection experiments targeting the dorsal striatum were visually selected based on proximity to our injection site. Four distinct sets of coordinates were included, and the projection volume (sum of segmented pixel counts) per brain region was averaged within each set of coordinates (Suppl. Fig. 5A). The four averages were subsequently combined (Suppl. Fig. 5A), and the resulting projection volume within each parcel of the fine-grained atlas was calculated to yield a structural connectivity vector. Only projections in the right hemisphere were considered, owing to ipsilateral bias of the anterograde experiments in the AMBCA. The values were log_10_-transformed and entries falling below -3.5 were set to zero to minimize false positives arising from tissue and small segmentation artifacts (*27*).

### Seed-based FC

Following confound correction, functional images were smoothed using a FWHM Gaussian kernel of 0.5 mm (*94*). The seed region was defined by thresholding the average structural-connectivity density map to encompass the region corresponding to the injection site in the dorsal striatum (Fig. 4A, Suppl. Fig. 5). Pearson correlation coefficients were computed between the mean time series of the seed region and the time series of every voxel in the brain and were converted to *z*-scores using Fisher *r*-to-*z* transformation for voxelwise analysis. Group-average seed-based FC maps were generated for each group by averaging the individual subject maps. The average connectivity within each parcel of the fine-grained atlas was calculated and used for subsequent correlation analysis.

### Atlas-based functional connectivity (FC)

Using the coarse atlas, the unsmoothed timeseries of all voxels within each parcel were averaged. Pairwise Pearson correlation coefficients between each pair of regions were calculated using nilearn (*94*) and then converted to *z*-scores using Fisher *r*-to-*z* transformation. Subject-level connectivity matrices were obtained by averaging across runs and subsequently used for group-level statistical inference.

### Independent component analysis (ICA)

To examine changes in correlation between different brain networks following PFF injection, ICA analysis was conducted. The smoothed data from all subjects was concatenated and FSL’s MELODIC (*95*) was used to conduct group-level independent component analysis (ICA). The data was decomposed into 30 networks using RABIES’ precomputed networks as priors (*29*). The priors were generated by running group-ICA with 30 components on the REST-AWK dataset (*29*). Five major networks were used for correlation analysis (Suppl. Fig. 8). The Pearson correlation coefficients were calculated between the timeseries of each pair of networks, converted to *z*-score and later used for group analysis.

### Paw movement tracking

While resting-state fMRI (rsfMRI) can reveal changes in FC and network organization, task-based fMRI provides useful insight about how specific networks are modulated during a defined task. Since PD is recognized as a movement disorder and PFF injection leads ultimately to severe motor deficits, we wanted to investigate how PFF injection affects the motor network during a motor task. Setting up a classical motor task for mice inside the scanner such as joystick movement is challenging. Instead, we leveraged the spontaneous paw movements during scanning in the awake mice.

Bilateral paw positions were tracked (Fig. 5A) across all 40 recording sessions (10 animals, 4 runs each) using DeepLabCut (v3.0.0rc13) with PyTorch 2.9.1+cu128 as a backend (*83*). Videos were acquired at 640 × 480 pixel resolution at 20 fps. Two body parts were annotated: the right paw and left paw (Fig. 5A). The DeepLabCut GUI was used to extract 20 frames per video across the full session duration using k-means-based sampling to maximize pose diversity. Frames from all 40 videos were manually labeled, and the combined dataset was used to create a single training set (95% training fraction, 5% held out for evaluation). A ResNet-50 convolutional neural network was trained for 200 epochs with a batch size of 8, using the default augmentation pipeline. Per-frame x, y coordinates and likelihood scores were exported as CSV files. Pose estimates with a likelihood below 0.6 were excluded from downstream analyses. The x and y coordinate timeseries for each paw were mean-centered and subjected to principal component analysis (PCA) with two components using scikit-learn (*96*). The first principal component (PC1) captured the dominant axis of paw motion across the session. Movement events were identified as peaks in the PC1 timeseries using scipy.signal.find_peaks (*97*). Each run typically contained multiple movement events, and the onset and duration of the events were used to construct a custom explanatory variable (EV) file for subsequent general linear model (GLM) analysis.

The GLM analysis was done using FSL (*98–100*). Paw movement events were convolved with a double-gamma hemodynamic response function (HRF) and its temporal derivative to produce the predicted BOLD response regressor for each paw separately. The spatially smoothed timeseries were then fit to the regressor in a voxel-wise manner using film_gls. Group-level inference was performed using a mixed-effects model (flameo) with a cluster-forming threshold (CFT) of Z > 2.1 and cluster-level correction at p < .05 using Gaussian random fields theory (GRFT). To extract the average timeseries, masks of four regions (Fig. 5C): the right somatomotor/somatosensory cortex, the lateral septal complex, the left somatomotor/somatosensory cortex, and left entorhinal area were derived from the atlas parcellations, the fitted timeseries from active voxels within each regional mask was averaged across voxels to yield a single regional timeseries per run. Stimulus-locked BOLD responses were extracted by windowing the average timeseries ±20 seconds around each paw movement event and averaging across events, runs, subjects, and finally across both groups. Group-mean BOLD responses were smoothed with a Gaussian filter (σ = 1 timepoint) prior to visualization (Fig. 5C). The average number of movements for each paw per session was calculated and compared between groups.

### Correlation analysis

To test how different structural and functional measures covary spatially with the pS129 field fraction in the α-synuclein PFF-injected animals, two complementary parcel-level spatial correlation analyses were conducted. In the first approach, each subject’s pS129 field fraction was spatially correlated with the corresponding parcel-wise W-score map and seed-based map, using Pearson *r* correlation coefficient, yielding a per-subject correlation coefficient for each measure that were then tested for significance across the group.

Second, group-averaged parcel-wise maps of pS129 field fraction, W-scores, and seed-based FC were constructed across PFF-injected animals, and Pearson correlation coefficients were computed between the pS129 map and each of the other measures. To account for the inherent spatial autocorrelation of brain maps — which inflates the effective degrees of freedom and produces spuriously low *p*-values — null distributions were generated using BrainSmash (*30*) for the W-score and the seed-based FC group-average maps, producing surrogate maps (n=1000) that preserve the spatial autocorrelation structure of each empirical map. The observed correlation coefficient was then compared against this null distribution to derive a corrected *p*-value.

Additionally, the group-averaged pS129 field fraction map in the right hemisphere was also correlated with the dorsal striatum structural connectivity vector (see above) using the same approach, with surrogate maps generated from the structural connectivity vector of the right hemisphere, to test whether the spatial distribution of pS129 pathology mirrors the structural connectivity profile of the injection site, consistent with a connectivity-based spreading model. Finally, the structural connectivity vector of the dorsal striatum within the right hemisphere was correlated with the group-averaged W-score map to test the correspondence between structural projections and morphological changes. The correlation of the dorsal striatum with itself was omitted, as the seed region’s self-projection is maximal by definition and represents a high-leverage point that is not informative about connectivity-dependent spread.

### Statistical analysis

For the DBM and seed-based FC voxelwise analysis, the relative Jacobian maps or the connectivity maps from each group were merged to form a 4D volume and an equivalent two-sample test was used to assess group differences. The average FC matrix from each subject (for atlas-based or ICA-based FC) was concatenated across subjects to form a group-level input and unpaired two-sample hypothesis testing was performed.

Group-level statistical comparisons were performed using non-parametric permutation testing, as implemented in the FSL Permutation Analysis of Linear Models (PALM) tool (*84*). Null distributions were generated from 10,000 permutations.

To compare pS129 field fraction between hemispheres, a paired Student’s t-test was conducted across each pair of ROIs for the coarse-atlas parcels and the results were corrected for multiple comparisons using Benjamini–Hochberg false discovery rate *(FDR; 101)*. Unpaired Student’s t-test was used to compare the average number of paw movements between groups.

Subject-wise correlations were evaluated using a one-sample Student’s t-test applied to Fisher *z*-transformed Pearson *r* coefficients. For group-average spatial correlations, statistical significance was assessed using spatially autocorrelation-preserving surrogate maps, which account for the non-independence of proximal brain regions and thereby control the false positive rate (*30*).

## Supplementary Figures

**Suppl. Fig. 1.**
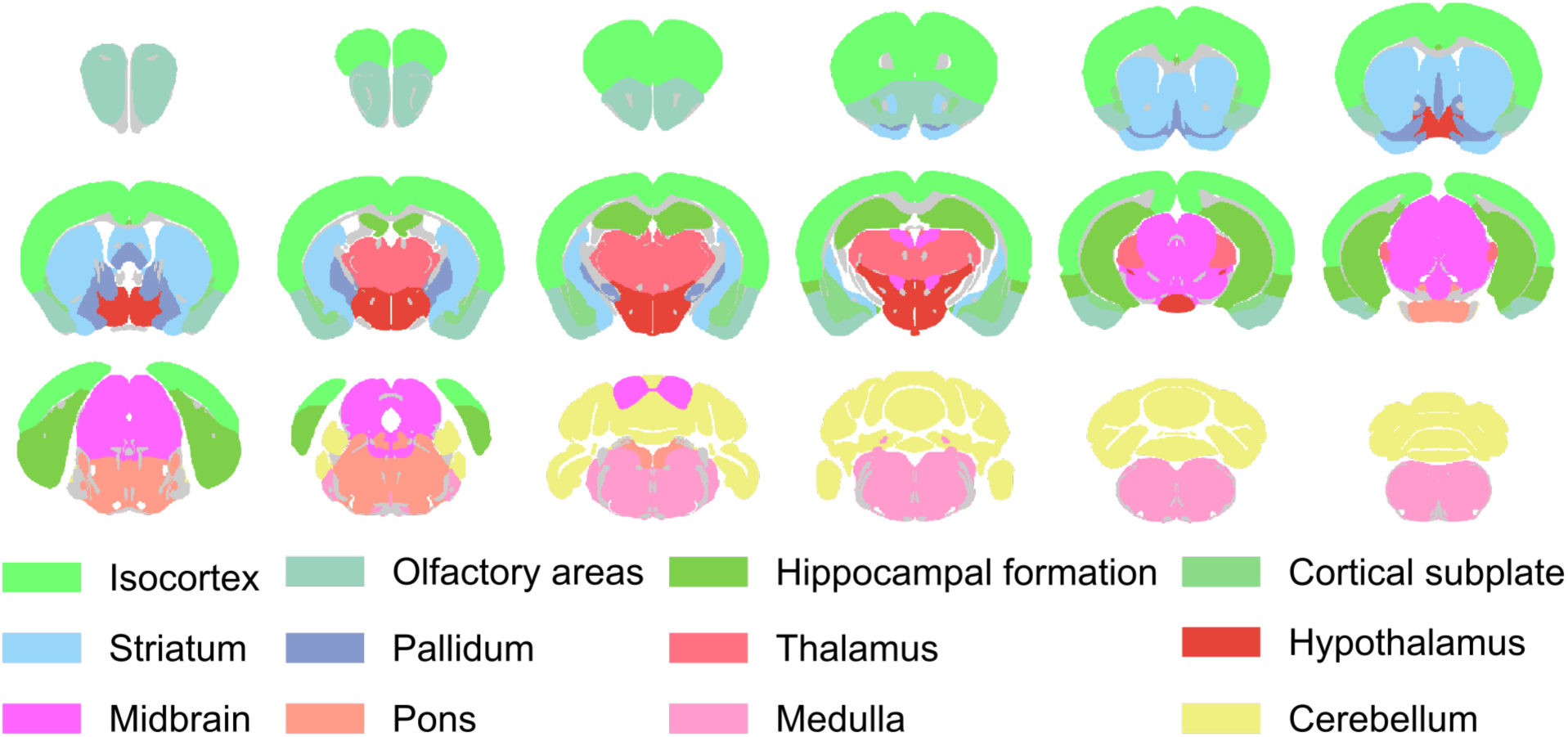
Coarse atlas parcellation. The parcellation comprises 12 regions per hemisphere, derived from the Allen Mouse Brain Common Coordinate Framework v3 (ABA-CCFv3).

**Suppl. Fig. 2.**
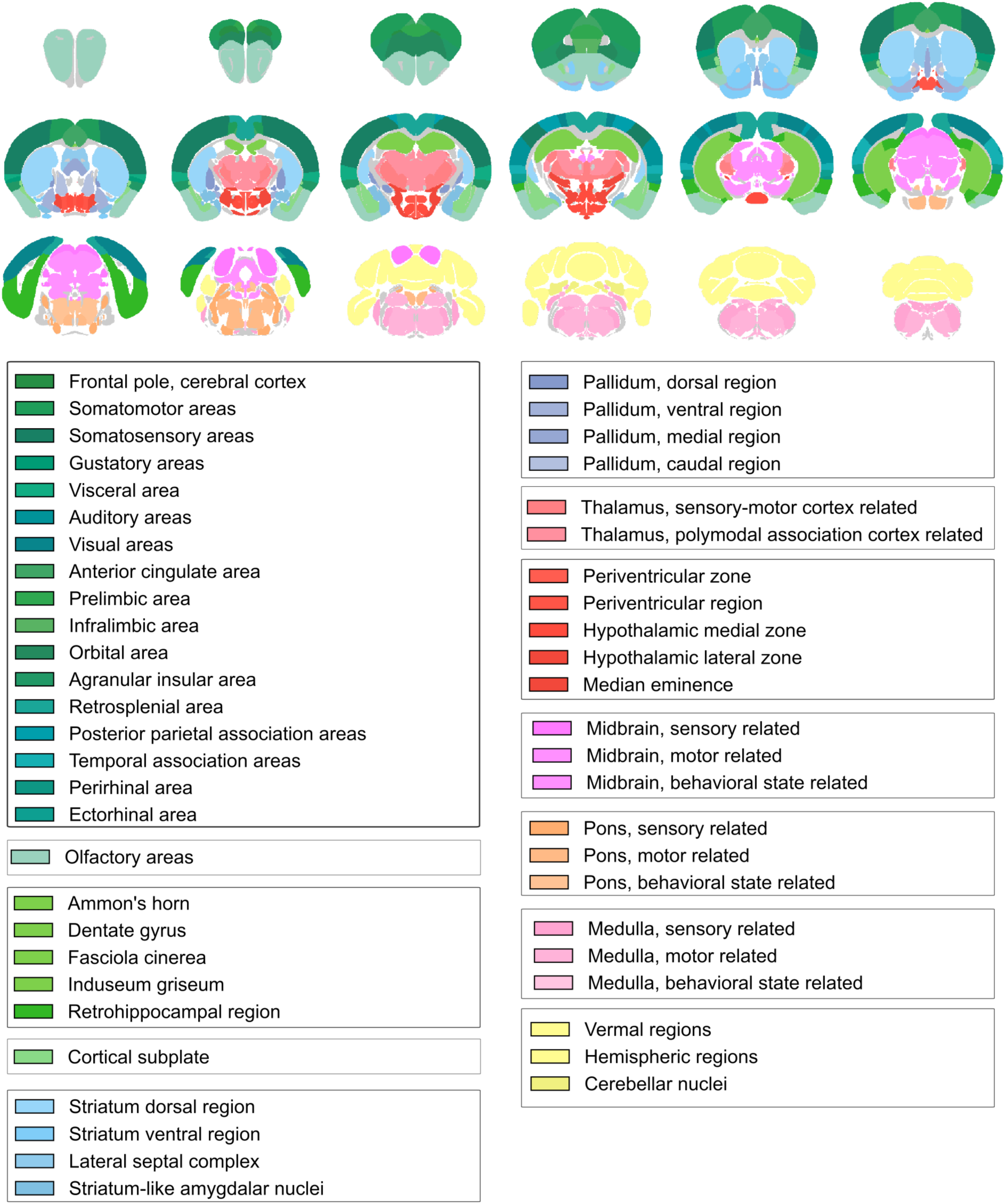
Fine atlas parcellation. The parcellation comprises 51 regions per hemisphere, derived from the Allen Mouse Brain Common Coordinate Framework v3 (ABA-CCFv3).

**Suppl. Fig 3.**
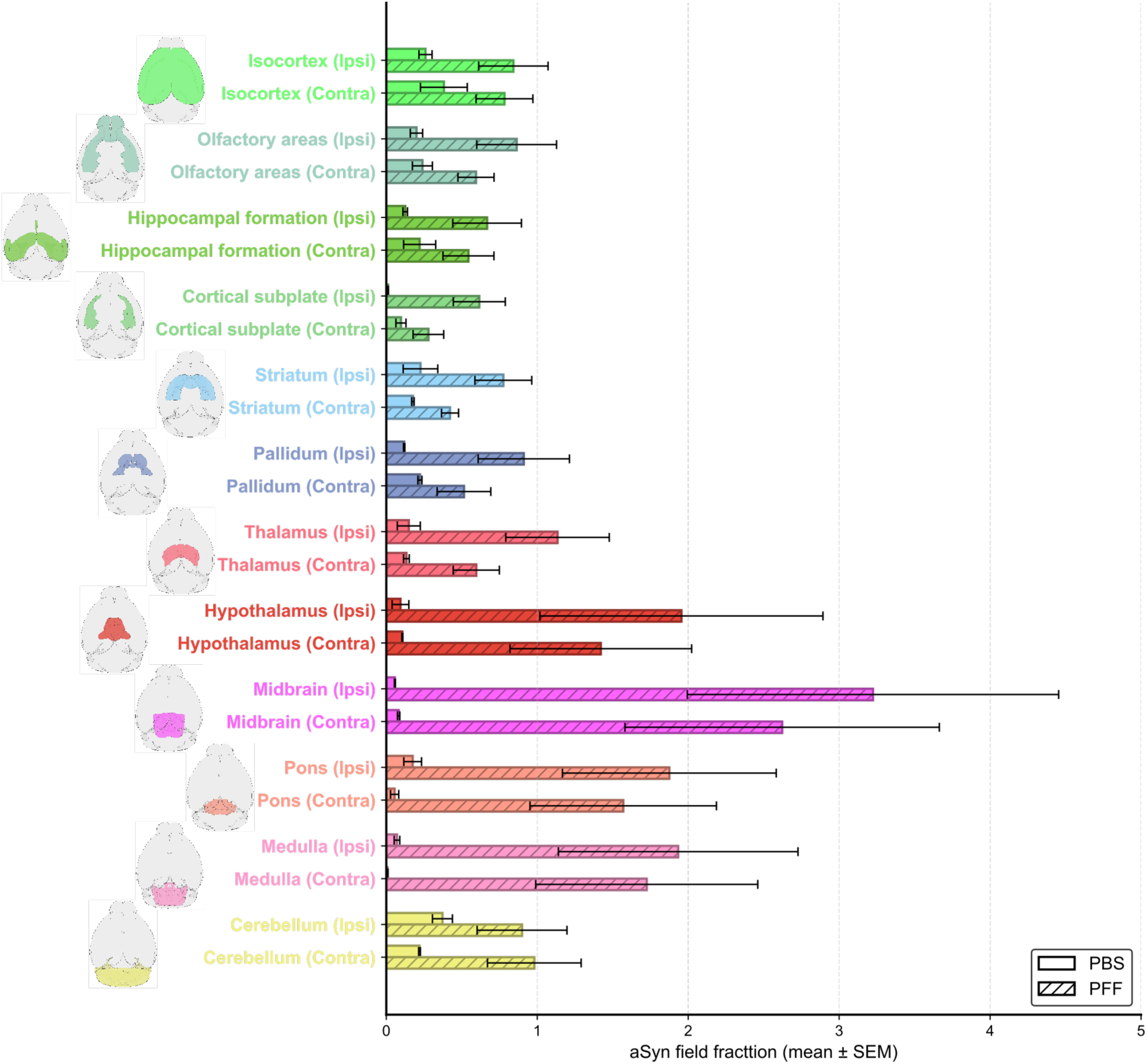
PS129 field fraction did not differ significantly between hemispheres. Quantification of pS129 across major brain structures in the ipsilateral and contralateral hemispheres (bars, mean ± SEM). **Abbreviations:** aSyn, α-synuclein; PBS, phosphate-buffered saline; PFF, preformed fibril; pS129, phosphorylated α-synuclein at S129; SEM, standard error of the mean.

**Suppl. Fig. 4.**
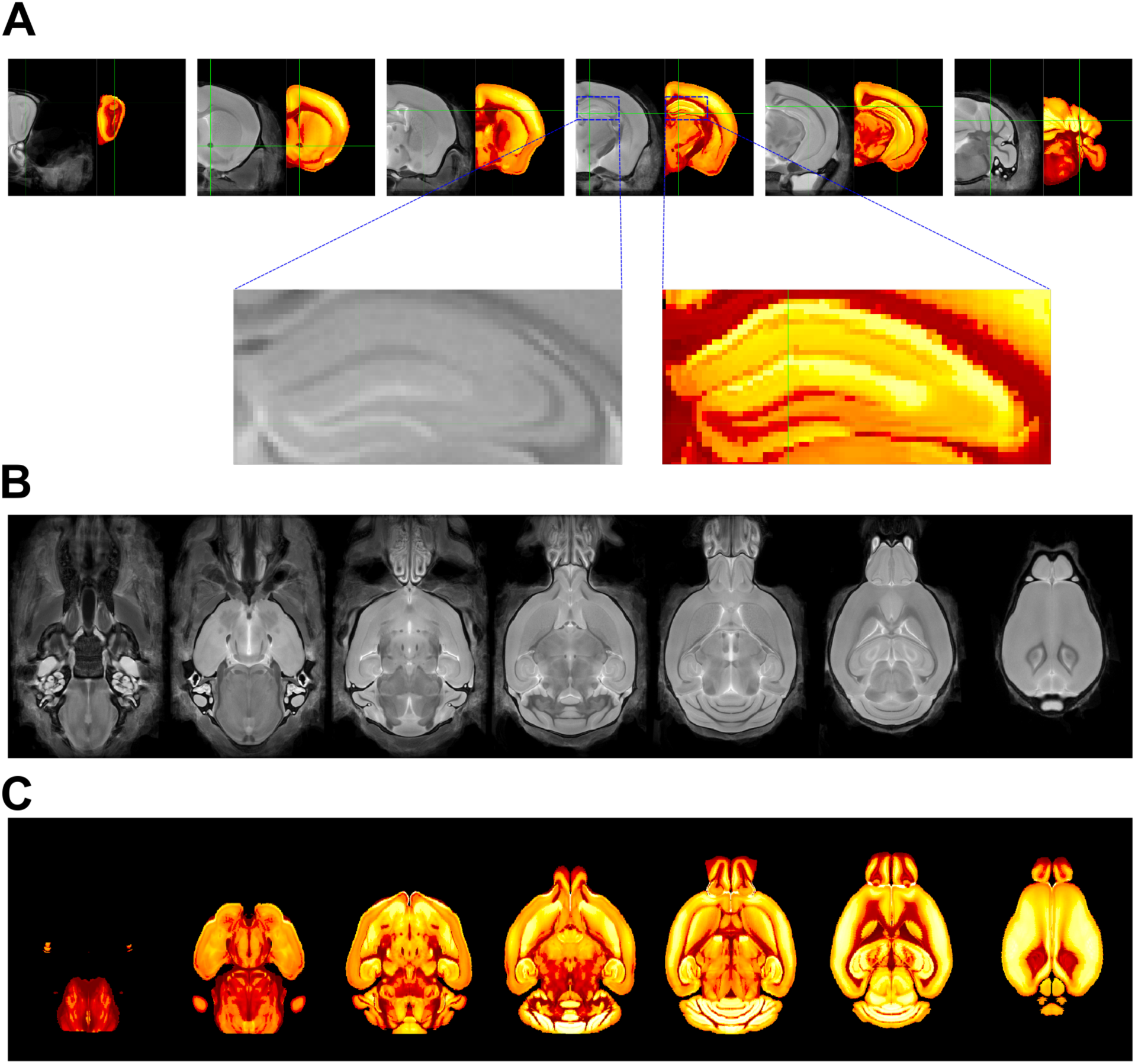
Registration of the ABA-CCFv3 anatomical template to the DSURQE template. **A)** Correspondence between the ABA-CCFv3 template, registered to DSURQE space, and the DSURQE template across representative coronal slices, with the hippocampus enlarged. **B)** Representative axial slices of the DSURQE template. **C)** Representative axial slices of the ABA-CCFv3 template. Abbreviations: ABA-CCFv3, Allen Mouse Brain Common Coordinate Framework v3.

**Suppl. Fig. 5.**
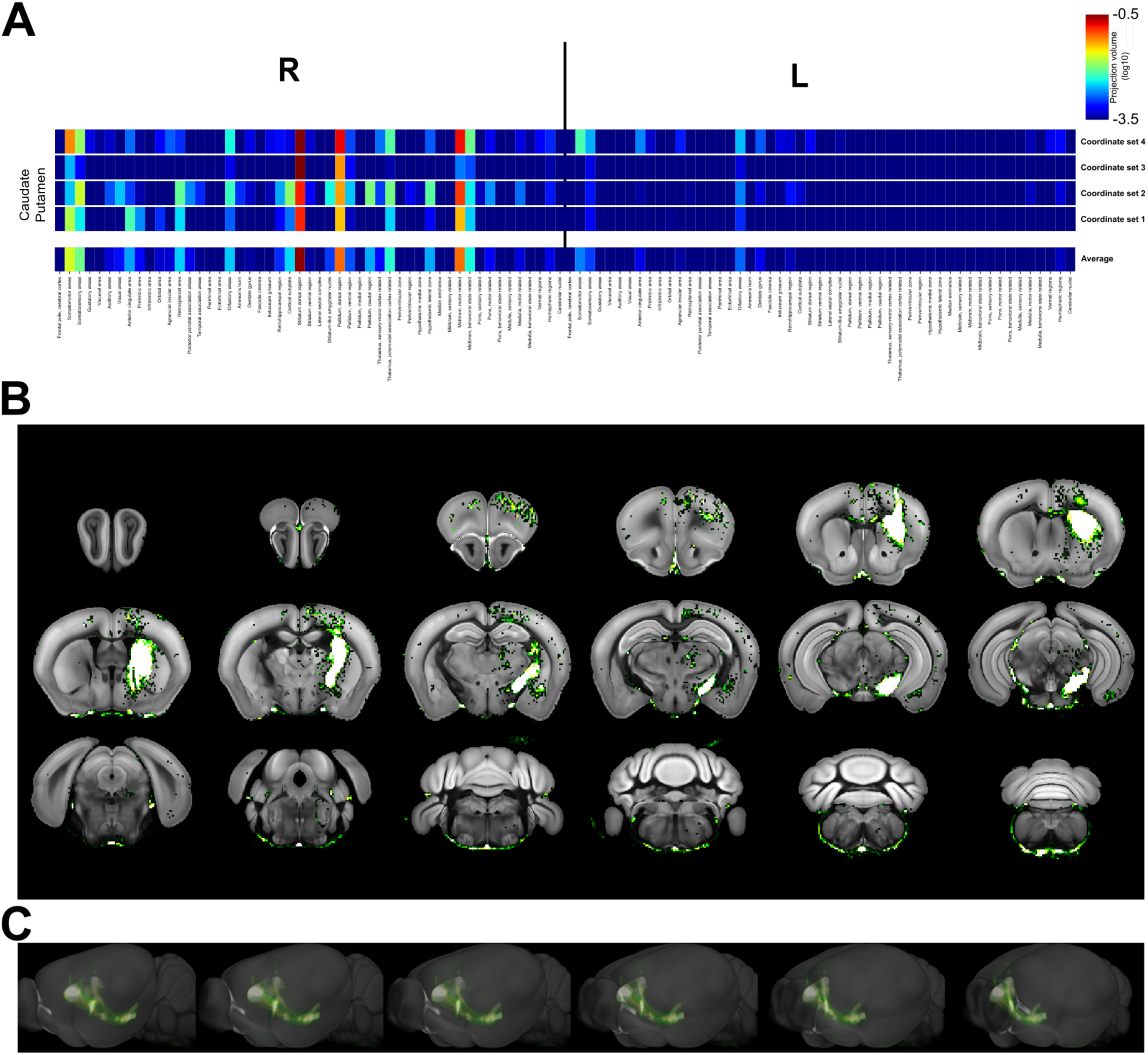
Structural connectivity of the dorsal striatum. **A)** Each row shows the average quantitative projection volume from the dorsal striatum to the rest of the brain across the right (ipsilateral) and left (contralateral) hemispheres for one set of injection coordinates; the final row is the average across all sets. Colormap represents log10-transformed values. **B)** Projection density map averaged across all coordinate sets. **C)** Thumbnails of 3D renderings of the average density map. Projection data are derived from the Allen Mouse Brain Connectivity Atlas (AMBCA) anterograde tracing. **Abbreviations:** AMBCA, Allen Mouse Brain Connectivity Atlas; CP, caudoputamen; L, left hemisphere; R, right hemisphere.

**Suppl. Fig. 6.**
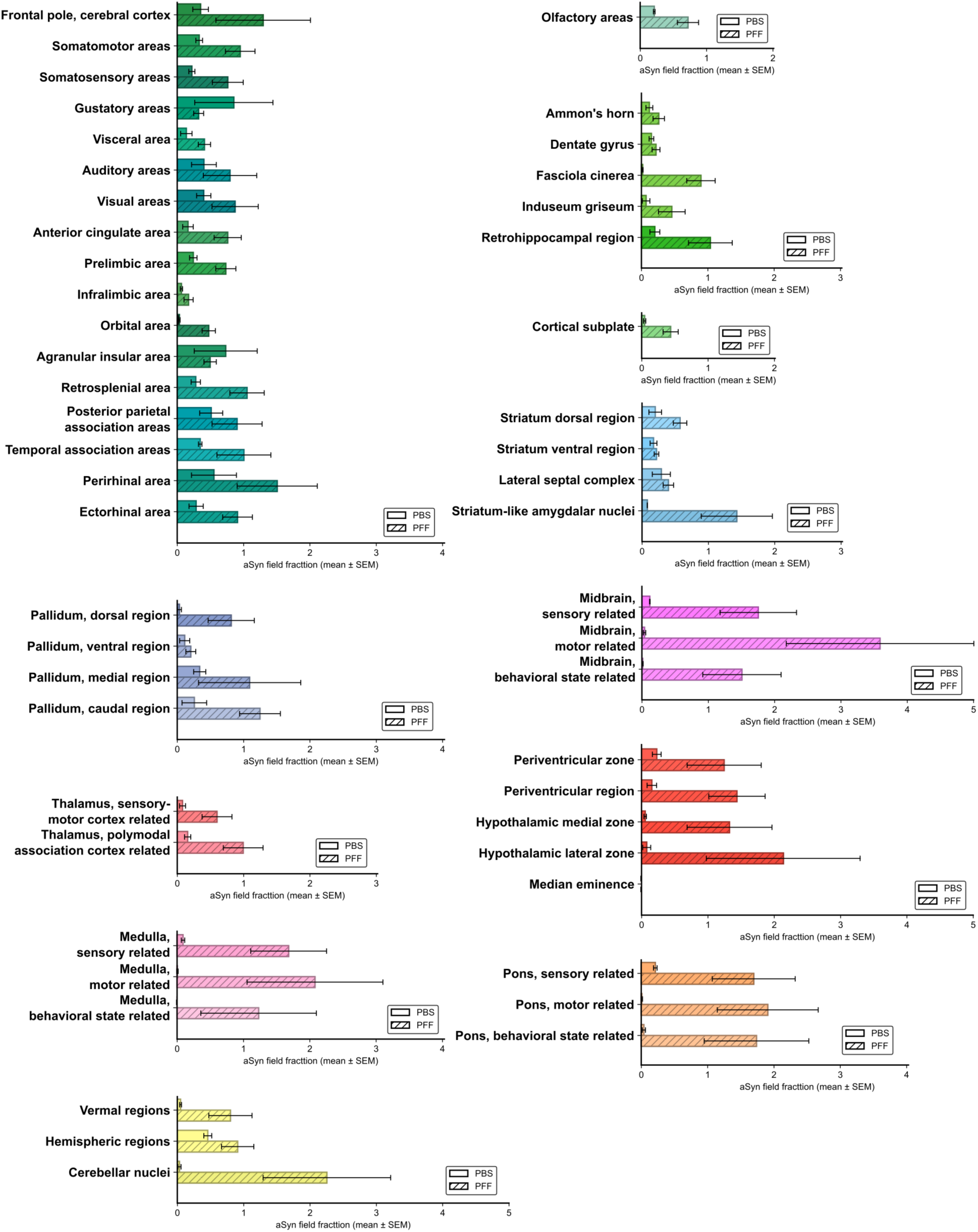
Quantification of pS129 field fraction across fine anatomical structures based on Allen Mouse Brain Common Coordinate Framework v3 (ABA-CCFv3) parcellations (, mean ± SEM). **Abbreviations:** aSyn, α-synuclein; PBS, phosphate-buffered saline; PFF, preformed fibril; pS129, phosphorylated α-synuclein at S129; SEM, standard error of the mean.

**Suppl. Fig. 7.**
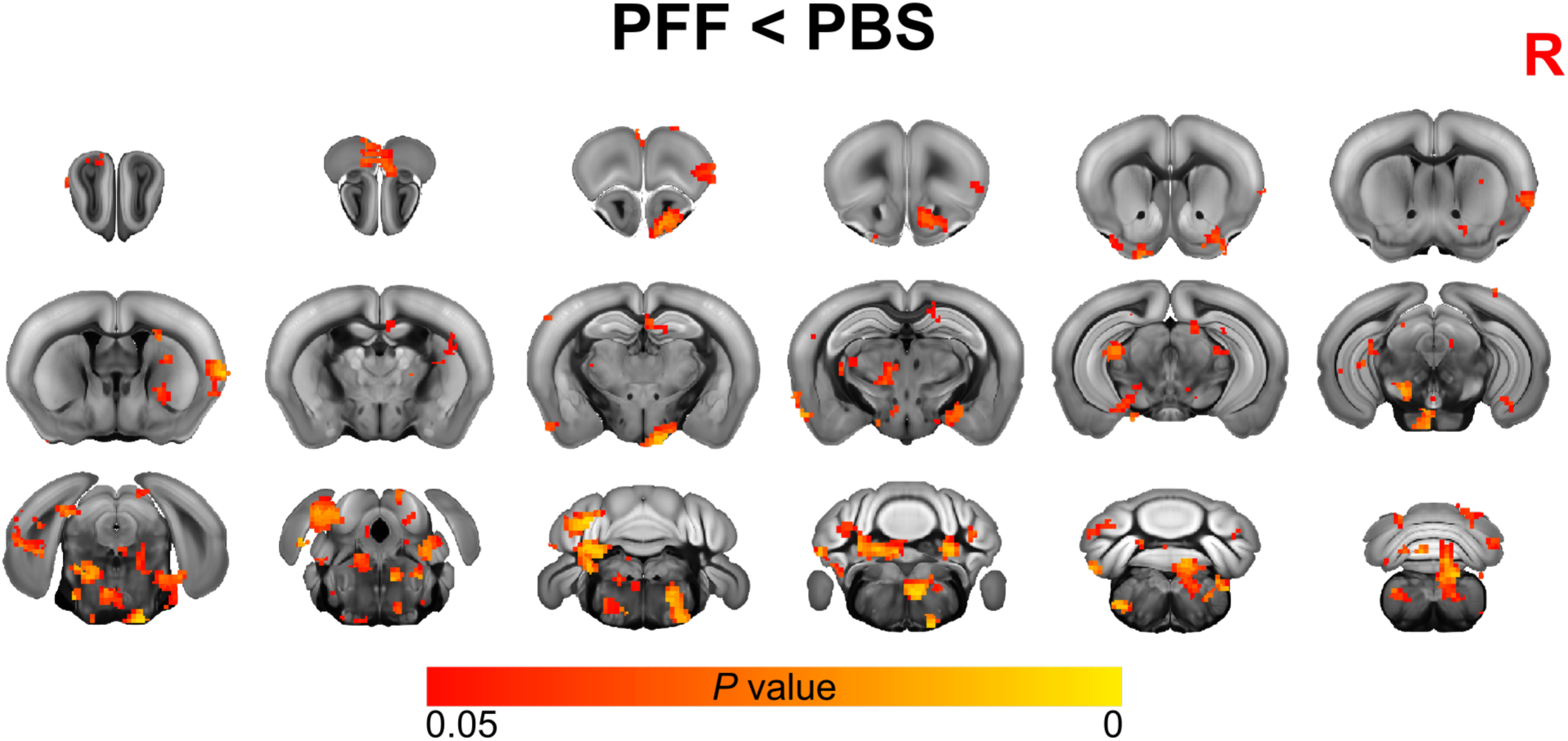
Statistically significant differences in seed-based connectivity between the dorsal striatum and the rest of the brain. Voxelwise map showing regions of decreased seed-based functional connectivity in PFF-injected compared with PBS-injected animals; the statistical map is overlaid on the ABA-CCFv3 template. **Abbreviations:** ABA-CCFv3, Allen Mouse Brain Common Coordinate Framework v3; FC, functional connectivity; PBS, phosphate-buffered saline; PFF, preformed fibril; R, right hemisphere.

**Suppl. Fig. 8.**
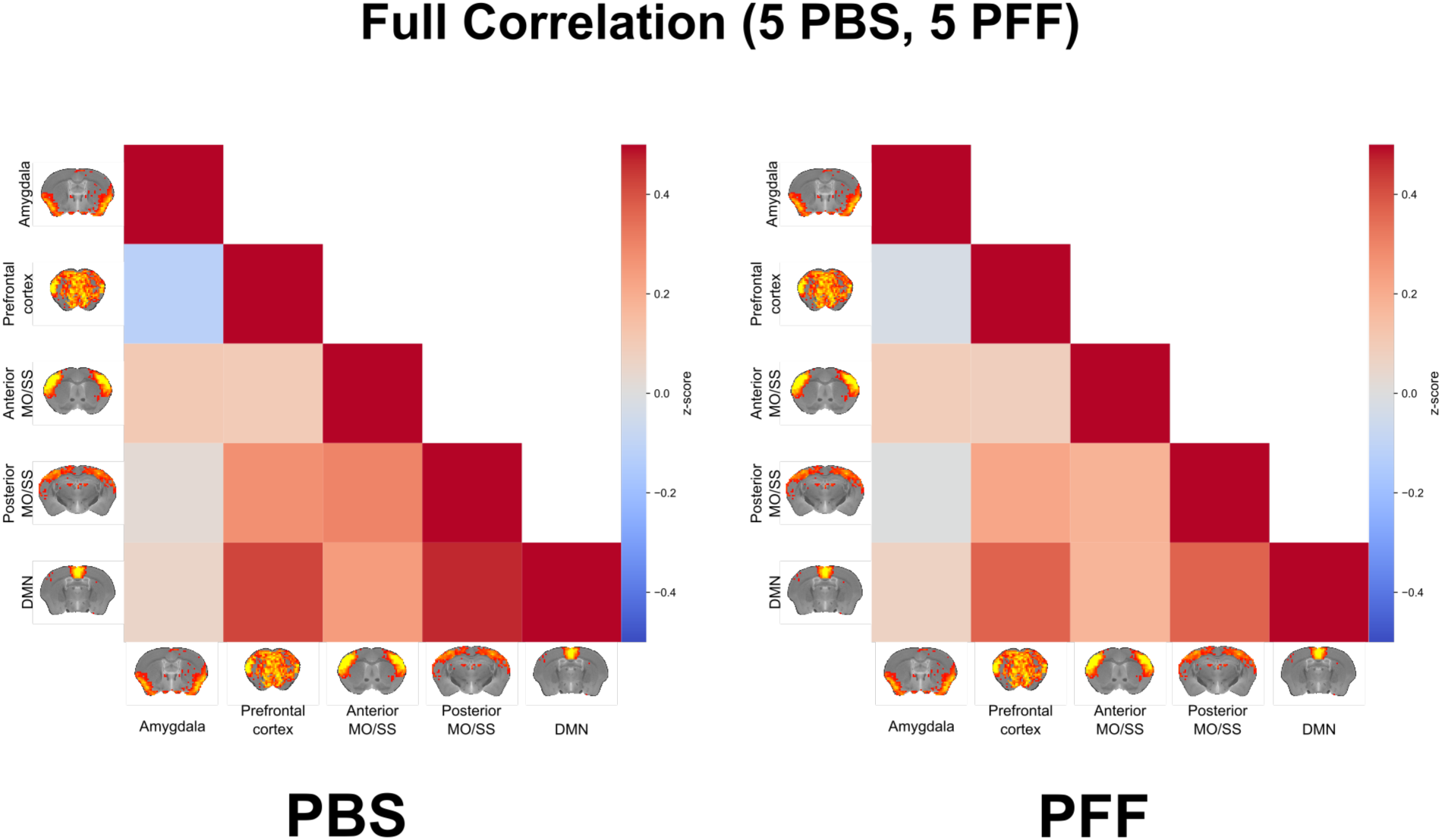
Functional connectivity between ICA networks did not show statistically significant differences between PFF- and PBS-injected animals. Average FC between ICA networks in PBS-injected animals (left) and PFF-injected animals (right). **Abbreviations:** FC, functional connectivity; DMN, default-mode network; ICA, independent component analysis; MO, somatomotor area; PBS, phosphate-buffered saline; PFF, preformed fibril; SS, somatosensory area.

**Suppl. Fig 9.**
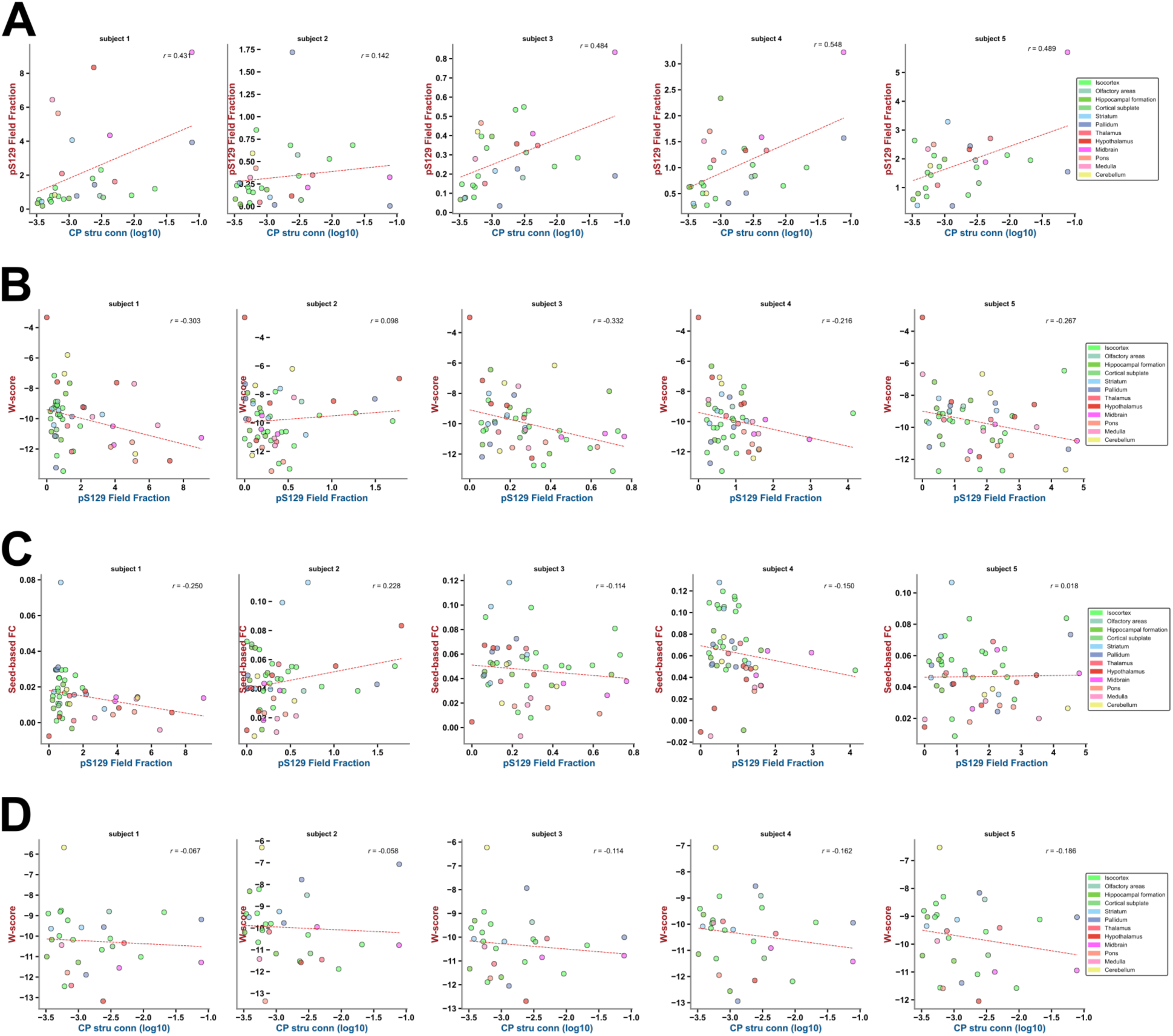
Per-subject correlations among pS129 field fraction, structural connectivity, atrophy, and functional connectivity. Each panel shows a scatter plot with Pearson’s correlation coefficient (r) for a single animal; rows correspond to the four correlations (A–D) and columns to the five individual subjects. **A)** Structural connectivity from the dorsal striatum (CP) versus pS129 field fraction within those regions, right hemisphere. **B)** Atrophy W-scores versus pS129 field fraction, averaged across both hemispheres. **C)** Seed-based functional connectivity versus pS129 field fraction, averaged across both hemispheres. **D)** Structural connectivity from the dorsal striatum (CP) versus atrophy W-scores within those regions, right hemisphere. Within each panel, dots of the same color denote substructures of a common major region; shading shows the confidence interval. Structural connectivity is derived from Allen Mouse Brain Connectivity Atlas anterograde tracing. **Abbreviations:** CP, caudoputamen; FC, functional connectivity; pS129, phosphorylated α-synuclein at S129; W-score, atrophy W-score.

